# High order expression dependencies finely resolve cryptic states and subtypes in single cell data

**DOI:** 10.1101/2023.12.18.572232

**Authors:** Abel Jansma, Yuelin Yao, Jareth Wolfe, Luigi Del Debbio, Sjoerd Beentjes, Chris P. Ponting, Ava Khamseh

## Abstract

Single cells are typically typed by clustering in reduced dimensional transcriptome space. Here we introduce Stator, a novel method, workflow and app that reveals cell types, subtypes and states without relying on local proximity of cells in gene expression space. Rather, Stator derives higher-order gene expression dependencies from a sparse gene-by-cell expression matrix. From these dependencies the method multiply labels the same single cell according to type, sub-type and state (activation, differentiation or cell cycle sub-phase). By applying the method to data from mouse embryonic brain, and human healthy or diseased liver, we show how Stator first recapitulates other methods’ cell type labels, and then reveals combinatorial gene expression markers of cell type, state, and disease at higher resolution. By allowing multiple state labels for single cells we reveal cell type fates of embryonic progenitor cells and liver cancer states associated with patient survival.

## 1. Introduction

To attribute disease to cell type, and molecular features of cells to disease state, we need to define and distinguish cell types, sub-types and states (Dann et al., 2023). The Human Cell Atlas (Regev et al., 2017) has taken a step in this direction by seeking definition of all human cell types and their molecular features, most often gene expression, within a multidimensional ‘cell space’ (Regev et al., 2017). Typing of cells is easiest when their lineages are well separated, and hardest when they are distinguished only by state (such as cell cycle phase, level of maturity, or response to stimulus) or spatial location. A common approach, whose theory is however problematic (Chari and Pachter, 2023), defines a cell type as a collection of cells that group more closely in gene expression space than other cells. This approach has yielded cell type definitions at relatively low-resolution, but requires additional analyses to begin resolving states within continuous trajectories of cell-state change (Ponting, 2019; Dann et al., 2022; Kotliar et al., 2019).

Cells adopt a continuum of states, representing cellular activities such as the cell cycle or responses to stimuli (Kotliar et al., 2019; Xia and Yanai, 2019). Labelling cells only by type thus does not finely resolve their dynamic behaviour such as during development or disease (Morris, 2019). Cell states are currently predicted by Principal Component Analysis (PCA) (Shalek et al., 2014; Steuerman et al., 2018), Independent Component Analysis (ICA) or Non-Negative Matrix Factorization (NMF) (Puram et al., 2017; Saunders et al., 2018). However, components or factors inferred by these algorithms may not faithfully or finely resolve cellular processes. States previously predicted by NMF among cancer cells, for example, include non-specific descriptors, for example ‘stress’, ‘metal response’, and ‘basal’ (Barkley et al., 2022).

To identify cell (sub)types and states at high resolution, we introduce Stator, which eschews cell clustering and instead defines states using the coordinated expression and non-expression of genes in single cells. Higher resolution is achieved by taking advantage of expression interactions at higher than pairwise order (3 ≤ *n* ≤ 7). Expression interactions that commonly co-occur in cells are gathered together as a single state label. The method yields biologically compelling labels of type, sub-type and state for cells in healthy and disease contexts without invoking concepts of expression space or pseudotime. These states are neither necessarily proximal in gene expression space nor necessarily categorical, thereby capturing the continuous nature of cell states and, in some cases, previously defined types. As with all cell state or (sub)type markers, Stator labels do not necessarily imply molecular mechanism. Rather, Stator reveals molecular and cellular heterogeneity and dynamics that would otherwise have been overlooked but can now be investigated experimentally. We show how Stator predicts, in a data-driven manner, sub-phases of the cell cycle, the future neuronal type and sub-type fate of mouse embryonic radial glial-like cell precursors, rare human endothelial cell sub-types and cycling transformed hepatocytes whose gene expression signatures are prognostic in liver cancer.

## 2. Results

Stator’s high-level workflow is illustrated in Fig. 1 with each step detailed in the Methods. Briefly, after performing standard Quality Control (QC) (Luecken and Theis, 2019), including doublet removal (Wolock et al., 2019), it initially restricts consideration to the most highly variable genes (HVG; often *N* = 1,000) (Wolf et al., 2018) followed by binarisation of gene expression. Binarisation does not substantially alter conclusions when analysing sparse data (Bouland et al., 2023, 2021; Qiu, 2020). Input to Stator is a cell (*M*) by binary gene expression (*G*) matrix (Fig. 1A). The model-free estimator of higher-order interactions (MFI) we introduced in (Beentjes and Khamseh, 2020) then estimates *n*-point interactions among *n* = 2, 3, …, 7 genes (Fig. 1B). Comparison between this estimator of dependence and other estimators such as correlation and mutual information is presented in Fig. S1 and Fig. S2 (Jansma, 2023a). In the next step (Fig. 1C), d-tuples are extracted. These are gene tuples significantly driving these interactions (Methods). This step is achieved by comparing expression of each tuple of genes in the MFI estimator to their expression under the null distribution of independence. Next, a new matrix of cell (*M*)-by-binary d-tuple (*K*) is created (Fig. 1D). Entries with 1 in the matrix indicate cells with that particular d-tuple gene expression combination; entries with 0 do not contain that given gene expression combination. Stator next performs hierarchical clustering of gene d-tuples based on these d-tuples’ co-occurrence in single cells (Fig. 1E). Crucially, this clustering takes place in a restricted space of d-tuples. Consequently, rather than a cell being placed at a single location in gene expression space, as is usual in scRNA-seq analyses, Stator allows for cells to adopt multiple biological states (Fig. 1F). We show below that a single cell can be thrice (or more) labelled (Fig. S3), for example as a radial glial-like precursor cell, as an astrocyte progenitor and a cell in G2/M cell cycle phases. Once groups of combinatorial gene signatures are identified, users can tune the modularity parameter that varies the granularity at which Stator states are resolved. Stator’s memory and run time are discussed in the Methods and Supplementary Material, 7.1, Fig. S4-S5.

**Figure 1:**
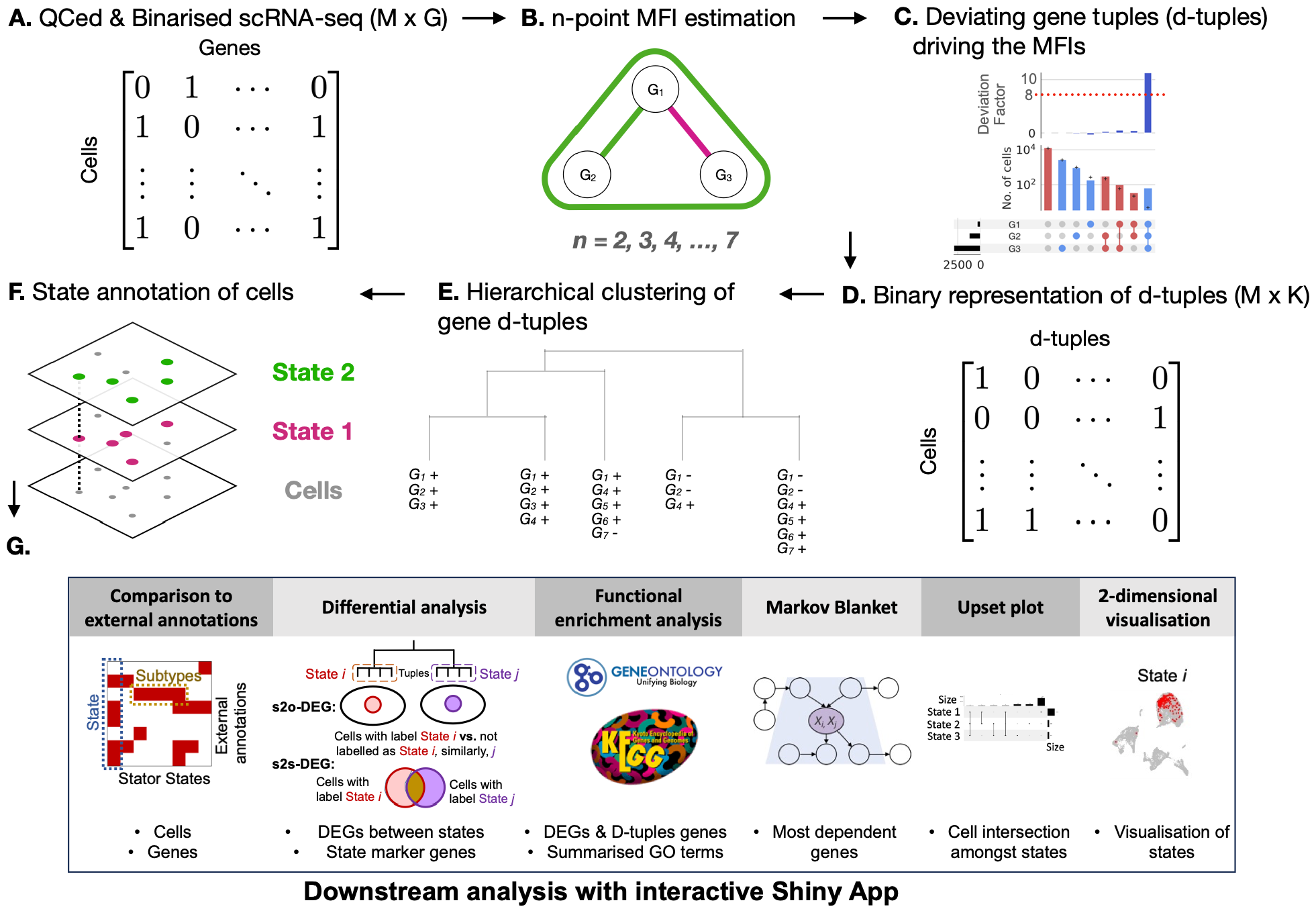
Workflow of Stator. Steps (A-E) are fully data-driven; steps (F,G) require biological interpretation. **(A)** A quality controlled, cell (*M*) by binarised gene expression (*G*) matrix is used as input. **(B)** *n*-point model-free interactions (MFI) are estimated (*n* = 2, …, 7) from the graph of conditional dependencies among the genes. Prior to this, the graph is inferred with an MCMC graph-optimisation algorithm on an initial structure obtained by the Peter–Clark causal discovery algorithm. The graph itself is not used to claim causation, rather to improve the statistical power of detecting *n*-point interactions among genes with strong inter-dependencies. **(C)** Tuples that are significantly deviating (default FDR *<* 0.05, log2FC=3) as compared to the null hypothesis of independence (interaction = 0) are extracted. These gene combinations are deviating tuples, or “d-tuples”. The significant tuple in this example is (*G*_1_, *G*_2_, *G*_3_) = (1, 1, 1) but d-tuples containing zero-values representing unexpressed genes are also possible. **(D)** A binary cell (*M*)-by-d-tuple (*K*) matrix is created. Entries with 1 indicate a cell containing a significant given tuple, in this example cells with (*G*_1_, *G*_2_, *G*_3_) = (1, 1, 1). Entries with a zero represent cells not containing the d-tuple. The matrix is created using all *K* significant interactions and corresponding d-tuples. **(E)** Hierarchical clustering of d-tuples (rather than cells) is performed to group any d-tuples that co-occur unusually often in single cells. The dendrogram is cut, by default, at a Dice-similarity that maximises the modularity score (Newman, 2006), but is adjustable. This procedure results in groups of d-tuples that can contain both the presence and absence of a gene’s expression. **(F)** At this stage, the user annotates and interprets the groups of d-tuple genes to infer cell states. Unlike clustering of cells, this procedure can result in cells that exist in multiple biological states simultaneously. **(G)** A Shiny App in R enables the user to compare Stator states against external annotations, such as other data-driven or expert annotations. Left: The horizontal box represents the significant enrichment of several Stator states in cells with a specific externally annotated cell type, demonstrating the existence of multiple cell subtypes that could be explored further. The vertical box represents Stator states spanning multiple externally annotated cell types, representing noncell-type restricted biological states, *e.g*., cell-cycle phases. The user can also choose to compare Stator state enrichment against biological conditions of an experiment. Stator’s Shiny App allows further integrative analyses, such as differential expression of Stator states or Gene Ontology term enrichment (Ashburner et al., 2000; Aleksander et al., 2023; Yu et al., 2012; Wu et al., 2021; Sayols, 2023).

Stator states are definable not just by d-tuple genes but also by other genes that are significantly differentially expressed relative to all other or one other state (Fig. 1G): these are state-to-other DEGs (s2o-DEGs, adjusted p-value *<* 0.05, |*Log*2*FC*| *>* 0.25) and state-to-state differentially expressed genes (s2s-DEGs, adjusted p-value *<* 0.05, |*Log*2*FC*| *>* 0.25), respectively (Fig. 1G). For this analysis, the unbinarised expression values of all genes (not only the 1,000 HVG) are considered using existing methods, *e*.*g*., Stuart et al. (2019). The app further permits Stator states to be queried for their enrichment in previously-derived annotations, such as experimental condition (healthy versus disease, different time points), or cell (sub)type or biological state labels.

To demonstrate Stator’s ability to identify cell types, subtypes and states we investigated three published scRNA-seq data sets in normal and disease contexts, in embryological or adult tissue, and in human or mouse. This first set contains astrocyte and neuron progenitors from mouse late embryonic (E18) brain (10x Genomics, 2017), chosen because this is the developmental stage when astrocytogenesis occurs and when cortical radial glial precursors (RPs) asymmetrically divide to generate neurons in the developing mouse cortex (Akdemir et al., 2020; Rubenstein and Campbell, 2020). The second is scRNA-seq data from human liver cells from control and disease (cirrhosis) donors (Ramachandran et al., 2019). Thirdly, we applied Stator to human liver cancer (hepatocellular carcinoma) cells (Barkley et al., 2022). Biological validity of a Stator state is provided when its d-tuple genes, s2o-DEGs and/or s2s-DEGs occur in a common cellular process and/or marker gene set. We start by showing how Stator identifies cells present in portions of cell cycle phases before then revealing cell subtypes and states that had hitherto not been inferred from these data sets.

### 2.1. Stator identifies states in seemingly homogeneous cells

We first applied Stator to 11,950 E18 mouse brain cells (Methods 4.6). These highly express canonical markers (*e*.*g*., *Slc1a3, Mt3* and *Mfge8* (Yuzwa et al., 2017)) of embryonic radial glial precursors (RPs), which later develop into astrocytes or neurons via intermediate progenitor (IP) cells. Upon clustering, these cells appear to be highly homogeneous without being separable into, for example, cells in cell cycle phases. Specifically, they could not be discriminated, using hierarchical clustering with significance quantification of clusters (Gao et al., 2022) on the original (unbinarised) expression space, beyond two significantly different (*p <*0.05 Bonferroni corrected, Methods) and robust clusters of 6,485 and 5,465 cells, respectively (Fig. S6).

By contrast, Stator predicted 25 States (Fig. 2A; Supplementary Table 1; Supplementary Table 2), with the optimal Dice similarity of 0.95. The majority (75.7%; *N* = 9, 044) of cells occupy one or more state, and 34.7% (4,151) of cells are unique to a single state (Fig. S3). Some states (*e*.*g*., #1 and #2) are localised in a PCA embedding, but most are not (*e*.*g*., #3 and #4). D-tuples in 7 states contained known cell cycle marker genes (Tirosh et al., 2016; Fischer et al., 2016): in the largest (#11; 2,168 cells), nearly all d-tuples contained one or more known G1/S phase markers’ genes (33 of 36 d-tuples; 92%): 23 d-tuples contained either *Gins2* or *Gmnn* or both (20, 14 or 11 d-tuples, respectively). For illustration, one d-tuple contains three known S-phase expressed genes (*Dnmt1, Hells Pcna*; (Giotti et al., 2018)) with their co-ordinated expression (*i*.*e*., 1-values) in 258 cells, which corresponds to *>* 6.5-fold deviation from the null hypothesis of independent expression (FDR *<* 0.01); 35 other d-tuples co-occur sufficiently with this d-tuple in these cells to be combined into this single Stator state (#11).

**Figure 2:**
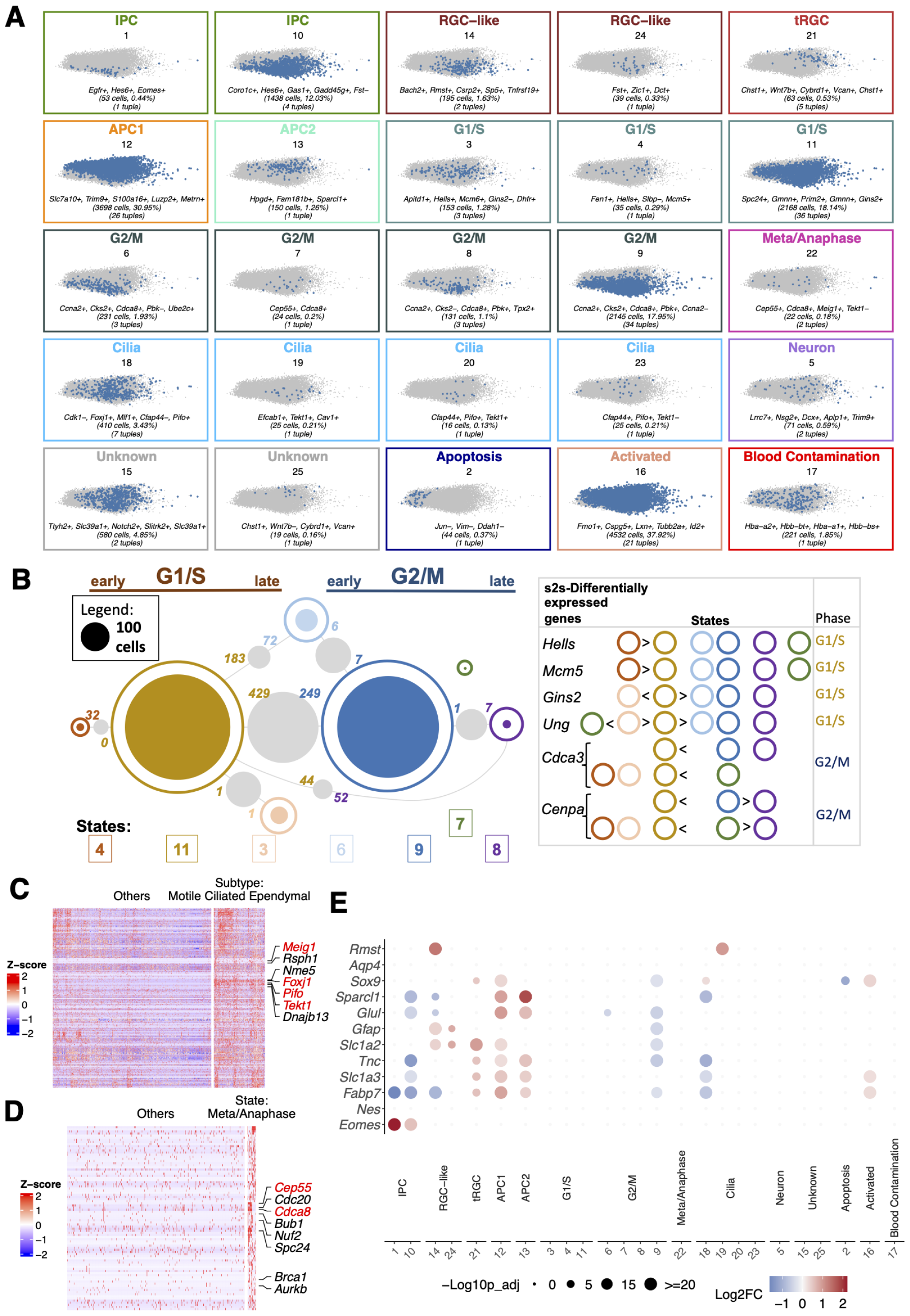
Stator identifies cell states in seemingly homogeneous mouse embryo radial glial cell-like precursor cells. **(A)** Stator identifies 25 signatures at maximum modularity. We have labelled 23 of 25 Stator states by performing differential gene expression analysis between cells in one state and all other cells, followed by gene enrichment (GO/KEGG) analyses. The significant differentially expressed genes were also compared with known gene markers of cell types and states. In such cases, we required at least 3 marker genes to be highly expressed. Every Stator state is highlighted in a PCA embedding of the unbinarised expression data, and annotated with the total number of cells and d-tuples it is composed of, as well as the five most common individual gene states across the states’ d-tuples. **(B)** Numbers of cells labelled with any one of 7 cell cycle states (# 3, 4, 6, 7, 8, 9 and 11); areas of circles are proportional to their number (see legend). Filled circles indicate numbers of cells labelled with only one of these single cell cycle states. Grey circles’ areas indicate numbers of cells labelled with two cell cycle states, those indicated by lines. Numbers of significantly differentially expressed genes between cell cycle state pairs (*i.e*., s2s- DEGs) are provided between the two states being compared; their colours refer to the state showing higher expression. State pairs with ≤ 25 cells are not shown. Right: s2s-DEGs are indicated by “*>*“ or “*<*“ symbols; for example, *Hells* mRNA expression is significantly higher in State 4 over States 11, 6, 9, 8 and 7. Early/late G1/S or G2/M cell cycle phase labels (top) were assigned using these mRNAs’ cell cycle phases known from high-throughput (top right; (Giotti et al., 2018)) and targeted experiments (*Ung* mRNA in late G1/S (Slupphaug et al., 1991) and *Cenpa* in G2 (Shelby et al., 1997)). **(C)** Heatmap of expression level (z-score) for genes up-regulated in state #18, versus other states, for cells in state #18 and a random selection of cells from other groups (*n* = 1,500). Z-scores are computed on a gene-by-gene basis by subtracting the mean and then dividing by the standard deviation throughout this study. Genes were ordered by hierarchical clustering. Upregulated genes are significantly involved in Cilium assembly (GO:0060271; *q* = 3 *×* 10^−11^). **(D)** Heatmap of expression level (z-score) for genes up-regulated in state #22, versus other states, for cells in state #22 and a random selection of cells from other groups (n=500). Genes were ordered by hierarchical clustering. Up-regulated genes reveal state of metaphase/anaphase. **(E)** Dot plot illustrating differential expression of astrocytogenesis marker genes across all Stator states. The size of the dots represents the -log_10_(Seurat p-val-adj) from differential expression testing between a state and all other states. Colour intensity represents the log_2_(FC) of gene expression.

To assess biological validity of these Stator predictions—whether they might indicate cell types, subtypes or states—we undertook differential gene expression analysis (Supplementary Table 3, Supplementary Table 4, Supplementary Table 5). The 7 states’ s2o-DEGs were predominantly cell cycle markers, confirming them as cell cycle states. Many s2s-DEGs were also cell cycle marker genes (Fig. 2B). Pairs of Stator states with s2s-DEGs are transcriptomically non-identical, even if they show some transcriptomic similarity, as expected for states located along a continuum. Note that pairs of states are concluded to be transcriptomically non-identical when they have significant s2s-DEGs (beyond the Stator state-defining d-tuple genes), thus contrasting these two states’ gene regulatory programs.

In the second largest prediction (state #9; 2,145 cells), all 34 d-tuples contained G2/M phase marker genes (Tirosh et al., 2016; Giotti et al., 2018; Fischer et al., 2016): 23 contained *Pbk*, 17 contained *Cenpa*, and 9 contained both. Stator predicted these states as G1/S (#11) and G2/M phases (#9), respectively, by their cells’ transcriptomes differing by 429 and 249 s2s-DEGs Fig. 2B) including for #11: G1/S phases’ marker genes (e.g. *Tuba1b, Rpa2, Mcm4, Tipin, Mcm2, Hat1, Rfc3* and *Rfc2*); and, for #9: G2/M phases’ marker genes (e.g. *H2afv, Arl6ip1, Stmn1, Ccdc34, Tacc3, Racgap1, Hmgb3, Calm3*, and *Cenpe*), all genes that did not contribute to d-tuple definition. As expected, cells co-labelled with both states #9 and #11 preferentially expressed G1/S marker genes (*Pclaf, Mcm6, Gins2* and *Gmnn*, for example) or G2/M markers (*Pbk, Cenpa, Ccnb2* and *Cdca3*, for example) compared with cells only labelled with state #9 or with #11, respectively. Stator thus not only identifies cells that are cycling, but further differentiates cells into G1/S versus G2/M cell cycle phases. This justified labelling our method’s predictions as “Stator states”.

Applying the same approach (comparing Stator states’ expressed d-tuple genes and s2s-DEGs with known cell cycle phase marker genes) identified five less-populated states (#3-4, #6-8) as additional cell cycle states, each transcriptionally non-equivalent with respect to states #11 (G1/S-phases) and/or #9 (G2/M-phases) and to each other, Fig. 2B. These five states’ s2s-DEGs again included marker genes for G1/S phases (states #3-4) or G2/M phases (states #6-8) relative to #11 (G1/S) and/or #9 (G2/M). In particular, 2 G1/S cell cycle phases’ marker genes (*Hells, Mcm5*) are significantly more highly expressed in cells in state #4 over #11, and indeed in states #3, 6, 9, 7 and 8; similarly, *Gins2* has higher expression in #11 than in #3, 6, 9 and 8, Fig. 2B.

Demanding that at least 3 s2o-DEGs are known markers of an annotation (Supplementary Tables 2-4), we labelled the other states as either Intermediate progenitor cells (IPC) (Ruan et al., 2021), radial glial cell-like cells (RGC-like) (Zheng et al., 2022), or astrocyte progenitor cells (APC) (Liu et al., 2022); or in the metaphase/anaphase of the cell cycle (significant enrichment of GO:0045841 (Ashburner et al., 2000), FDR*<* 0.05) or apoptosing or activated cells (expressing mitochondrial genome genes or intermediate early genes or activation markers (Lacar et al., 2016), respectively); or blood cell contaminants that highly expressed not just globin genes (Biagioli et al., 2009) but also *Alas2*, an erythroid-specific gene (Fig. 2). More specifically, from differential expression of s2s-DEGs *Sparc* and *Sparcl1* (Supplementary Table 6), states #12 and #13 appear to label two known astrocyte progenitor cell types (Liu et al., 2022), and state #21 is associated with higher expression of truncated radial glial cell markers (*Anxa2, Cryab*, and *Tmem47* (Yang et al., 2022)) relative to APC1 cells (state #12). We illustrate raw gene expression differences defining states #18 (Cilia) and #22 (Metaphase/Anaphase) in Fig. 2C, D. In Fig. 2E, we show how expression of the few established markers of precursor and intermediate cell states (Akdemir et al., 2020; Götz et al., 2015) varies across the 25 Stator states.

Stator was also applied to a second subset (*N* = 11,950) of the E18 RPs, independent of the first, replicating APC1, APC2, IPC, and RGC-like states, multiple G1/S and G2/M cell cycle phases’ states, and activated and blood contamination states (Fig. S7, Supplementary Table 7, Supplementary Table 8, Supplementary Table 9, Fig. S3B).

### 2.2. Cell cycle states in embryonic neurons and RPs

We next showed that Stator can also identify cells in G1/S or G2/M phases within an admixture of two cell types, neurons and RPs (*n* = 13,605 and 5,395), from a single E18 mouse brain (10x Genomics, 2017) (Methods). In all, Stator predicted 110 states from these combined cells (Supplementary Table 10, Supplementary Table 2), of which 57 were common to both neurons and RPs, 34 were highly-specific (≥ 99%) to neurons, and 19 to RPs. The median number of predicted states for a cell was 3 (Fig. S3C). Among 12 cell cycle Stator predictions were G1/S (#51), S/G2 (#55), early G2/M (#58) and late G2/M (#59) states (Fig. 3, Fig. S8, Supplementary Tables 10-12). RP cells in G1/S cell cycle phases (States #51 and #55) had not previously been detected in this embryonic stage by cell cycle classification (Yuzwa et al., 2017). An additional state (#57) involving cells that were predominantly labelled neurons (86%) showed multiple s2s-DEG markers for both newborn neurons (*e*.*g*., *Dcx, Tubb3, Gad2, Stmn2*) and G2/M phases (*e*.*g*., *Cenpa* and *Cdca3*) and hence is likely a post-G0 phase neuron state (Supplementary Tables 10-12).

**Figure 3:**
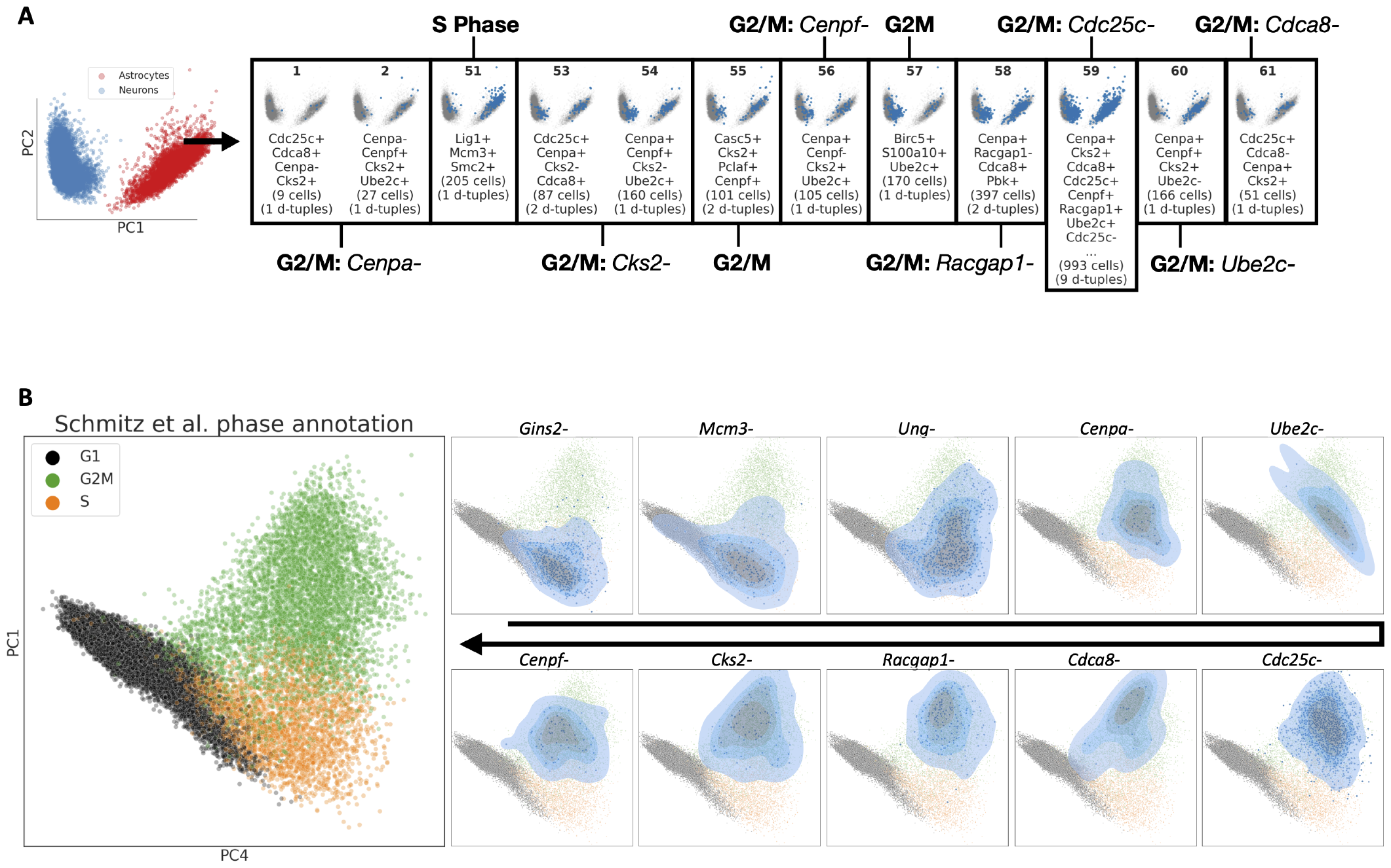
Stator identifies states present in two different cell types. **(A)** Stator states labelling both developmental neurons and radial glial cell precursors (RPs) from an E18 mouse brain data set (10x Genomics, 2017). Stator identifies 12 states that include one or more cell cycle phase gene markers as d-tuple genes, and that delocalise throughout expression space. State #51’s d-tuple genes (*Lig1+, Mcm3+, Smc2+*) encode proteins active in S phase, while the remaining 11 express known markers of G2/M phase (Riba et al., 2022; Tirosh et al., 2016; Giotti et al., 2018). It is notable that many G2/M states are defined by an absence of expression of d-tuple genes that are nevertheless known G2/M marker genes (*Cenpa, Cks2, Cenpf, Racgap1, Cdc25c, Ube2c* and *Cdca8*). **(B)** Left: Externally-derived cell cycle annotations of a mouse brain data set sourced from five different experiments (Schmitz et al., 2022), with cells from the 10X Genomics E18 mouse data set GSE93421 removed, mostly separate along principal components 1 and 4. Right: Cells and embedding as left, but marked by the expression of all but one of the marker genes. Different intensities of blue represent ‘densities’ of cells in the 2D embedding. Note that since embeddings can distort distances, densities cannot be directly interpreted so no legend or axis is shown. Each state contains expression of known cell cycle markers as well as a single non-expressed cell cycle marker gene (indicated above each box), predicted to be a combinatorial marker by Stator in RPs and/or neurons. These gene combinations thus demarcate cell cycle sub-phases, and a suggested ordering is shown here.

Eight G2/M states contained *minus* gene markers (*i*.*e*., those without expression evidence) that are nonetheless known markers of G2/M phases: *Cenpa* for States #1 and #2, *Cks2* #53 and #54, *Cenpf* #56, *Racgap1* #58, *Ube2c* #60, or *Cdca8* #61. We found similar combinatorial markers for the cell cycle in RPs, which included S-phase states with a lack of expression of the S-phase markers *Ung, Mcm3, Gins2*. To investigate whether these states demarcate portions (sub-phases) of G2/M cell cycle phases, we highlighted cells from an external data set along a cell cycle projection that expressed all but one G2/M or S-phase cell cycle marker genes, specifically the *minus* gene marker (Fig. 3, Fig. S8). This illustrated that cells in these Stator states differentially occupied parts of the cell cycle continuum, consistent with cell cycle sub-phases. For example, Stator states differentiated between early G2/M (d-tuples with an absence of *Ube2c* or *Cenpa* expression, *i*.*e*., *Ube2c-*, or *Cenpa-*), early-or mid-G2/M (*Cdc25c-* or *Racgap1-*), or mid-to-late G2/M (*Cks2-* or *Cdca8-* or *Cenpf-*) or early S-phase (*Gins2-*), mid-S-phase (*Mcm3-*), and late S-phase (*Ung-*) (Fig. 3B). Rather than single genes, it is the combinatorial gene expression pattern that provides high resolution of cell states. This is because populations of cells defined only by the expression of various combinations of cell cycle marker genes, without requiring that the *minus* gene is unexpressed, are not localised to a cell cycle (sub)-phase (Fig. S9).

Having successfully identified cell cycle sub-phases for RPs and for a combined RP and neuron data set, we next used Stator to identify additional cell states within the combined data set. Embryonic RPs were previously described as homogeneous at E17.5 (Yuzwa et al., 2017). By contrast, 47 Stator states could be labelled as RPs either because their d-tuple genes were embryonic RP markers (Yuzwa et al., 2017) or else they significantly more highly expressed such genes over all other states (Supplementary Tables 10-11, Supplementary Table 2). Of these 47 RP states, 21 were transcriptionally heterogeneous owing to their d-tuples including a *minus* gene marker, such as *Hes5* (states #26-27), *Qk* (#29), and *Pax6* (#34), each of which is involved in neural progenitor cell fate choice ((Imayoshi and Kageyama, 2014; Takeuchi et al., 2020; Ericson et al., 1997)).

These RP states were transcriptionally heterogeneous (Fig. S10, Supplementary Tables 13): (i) 13 RP states yielded large number of s2s-DEGs, compared with the most populous RP state (#44); (ii) 3 states (#13, #36 and #39) showed significantly lower expression of 7 core RP genes, *Mt3, Phgdh, Slc1a3, Ddah1, Aldoc, Vim*, and *Fabp7* (Yuzwa et al., 2017) than state #44; (iii) 15 states contained G2/M cell cycle phases’ marker genes among their s2s-DEGs relative to state #40; (iv) and 15 states yielded large ribosomal subunit genes as s2s-DEGs with state #40, a transcriptional signature of embryonic RP reactivation to become activated neural stem cells (Borrett et al., 2022; Dulken et al., 2017).

Thirty-four RP states had neuronal marker genes among their s2s-DEGs with states #40 or #44 (Supplementary Tables 12-13), consistent with these embryonic RPs having a future neuronal fate. Seventeen states co-express *Ascl1* and *Neurog2* (often with *Gadd45g*, a transcriptional target of ASCL1), two genes that are expressed in more mature cells in a mutually-exclusive manner (Parras et al., 2002). These states thus likely label early neural progenitor cells that have yet to acquire their GABAergic (*Ascl1*) or glutamatergic (*Neurog2*) neuronal fate in the forebrain.

### 2.3. Neuronal states

For our final analysis of embryonic mouse brain cells, we analysed two disjoint subsets each containing 19,000 mouse E18 neurons. As the modularity was maximised at Dice similarity of 0.97 and 0.91, respectively, we applied a mean similarity of 0.94, resulting in 29 states in each (Supplementary Tables 14-15), allowing us to compare the disjoint subsets at equivalent resolution.

This number of Stator states was five-fold more than the 4 pairwise significantly distinct clusters found by hierarchical clustering in expression space for the first disjoint dataset (Fig. S11). Stator successfully distinguished striatal medium spiny neurons (MSN) from interneurons by known marker genes’ expression (*e*.*g*., *Ngef, Nrxn1, Pou3f1, Tshz2* (Arlotta et al., 2008; Fuccillo et al., 2015; Su-Feher et al., 2022), versus *Arx, Epha5, Lhx6, Prox1* (Poirier et al., 2004; Li et al., 2022; Miyoshi et al., 2015), Fig. 4A, Supplementary Table 2, Supplementary Tables 16-17). It further separated MSNs into their two known sub-types, Direct or Indirect pathway cells (Cui et al., 2013; Cirnaru et al., 2021) via markers: Direct: *Ebf1, Foxp1, Isl1, Nrxn1, Zfhx3*, and *Zfp503* (Li et al., 2022; Fuccillo et al., 2015; Zhang et al., 2019; Shang et al., 2022; Precious et al., 2016) and Indirect: *Adora2, Ebf1, Gucy1a3* and *Gucy1b3* (Li et al., 2022) (Fig. 4B), and separated interneurons into *Htr3a* and/or *Npy* expressing subtypes (Tremblay et al., 2016) (Supplementary Table 2, Supplementary Tables 16-17). Three RP-like states were additionally detected (Fig. 4C). States could be further labelled as early or late via markers of neuronal maturation (Rubenstein and Campbell, 2020), specifically the temporal sequence of expression of *Dlx2, Dlx1, Dlx6os1* and *Dlx6* genes, and the later expression of MSN or interneuron markers (Fig. 4D, (Liu et al., 1997)).

**Figure 4:**
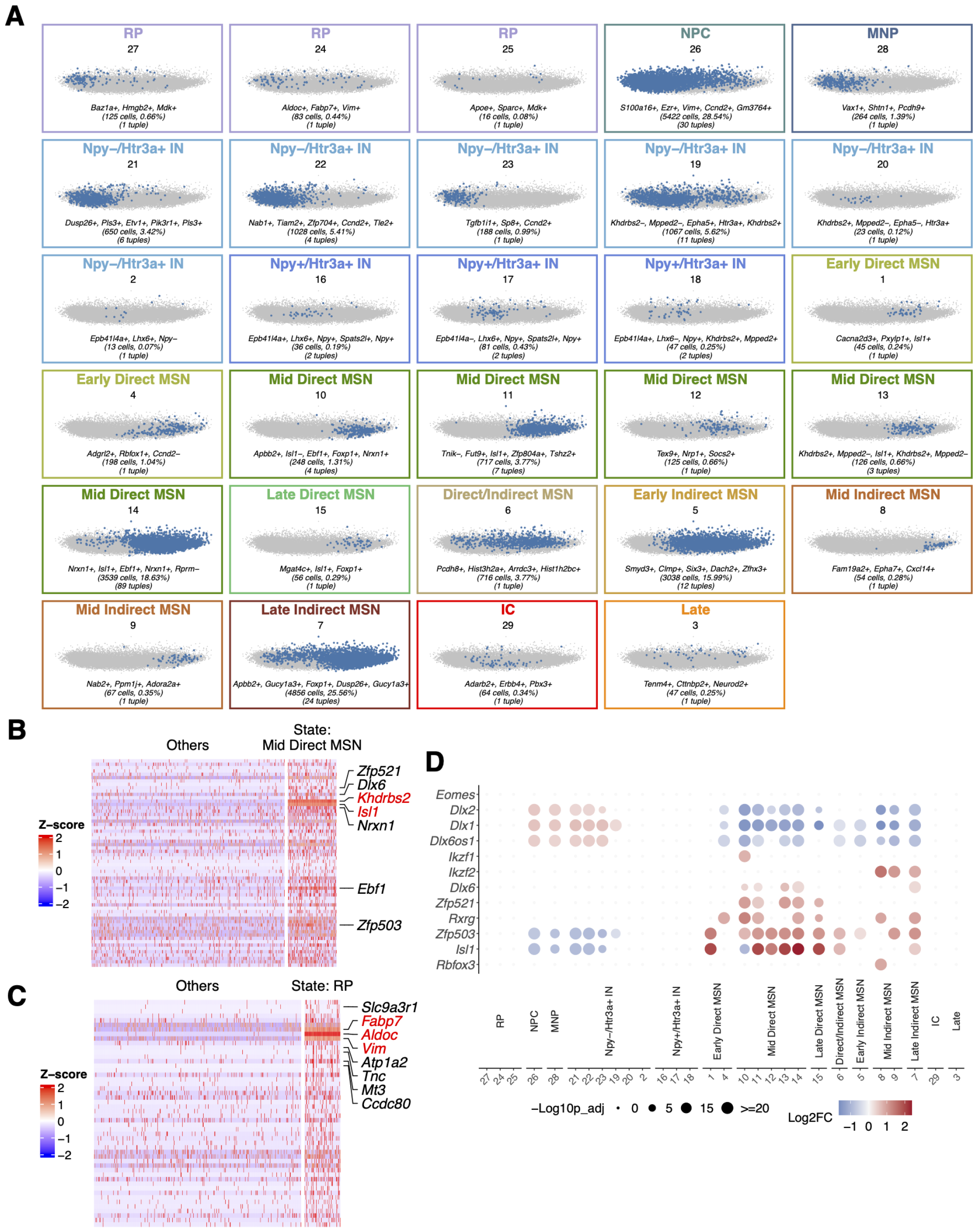
Stator identified states for developmental neurons. **(A)** Stator states for 19,000 E18 mouse cells, previously annotated as neurons. States were labelled by matching s2sand s2o- DEGs with literature gene markers, as before (Fig. 2). Abbreviations: IC, intercalated cells of amygdala (*Erbb4* +, *Tshz1* +, *Foxp2* +, *Pbx3* + (Peters et al., 2023; Kuerbitz et al., 2018); IN, interneurons; Late, late born neurons; MNP, migratory neuronal precursors (*Vax1* +, *Shtn1* +, *Pcdh9* +, *Tiam2* + (Coré et al., 2020; Sapir et al., 2013; Asahina et al., 2012; Kawauchi et al., 2003); MSN, medium spiny neurons; NPC, neural precursor cells; RP, radial glial cell precursors. **(B)** Heatmap of expression level (z-score) for genes up-regulated in state #13: Mid Direct MSN, versus other states, for cells in state #13 and a random selection of *n* = 500 cells from other groups. **(C)** Heatmap of expression level (z-score) for genes up-regulated in state #24, versus other states, for cells in state #24 and a random selection of *n* = 500 cells from other groups. **(D)** Dot plot illustrating differential expression of neurogenesis marker genes (Rubenstein and Campbell, 2020) across all Stator states. The size of the dots represents the -log_10_(Seurat p-val-adj) from differential expression testing between a state and all other states. Colour intensity represents the log_2_(FC) of gene expression.

Increasing the resolution of Stator state identification can resolve multiple constituent biological states. At a Dice similarity of 0.94, Stator’s state # 26 labelled neural precursor cells, as evidenced by high expression of *Zeb2, Mdk, Ctnna1, Arx*, and *Prox1*. Nevertheless, this state was found to be a composite of three component sub-states largely following the branching order of co-occurring d-tuples (see Fig. 1 D and E; Methods). From their d-tuple genes, these sub-states are readily distinguished as labelling G2/M cell cycle phases, neural stem cells and newborn neuronal precursors, respectively (Fig.S12).

Stator states representing the same neuronal subtypes (*e*.*g*., interneurons, direct or indirect MSNs and late born neurons) for the second disjoint dataset are shown in (Fig. S13, Supplementary Tables 18-19).

### 2.4. Stator resolves cell (sub)types in human liver disease at higher resolution

To demonstrate application of Stator in a human disease context, we analysed 20,000 cells from patients with uninjured or cirrhotic livers. These cells had previously been annotated as one of 12 types (Ramachandran et al., 2019). Stator identified 53 states (Supplementary Table 20), 28 that were differentially enriched between cirrhotic and uninjured liver sample cells (Fig. 5A). Enrichment of these states showed that Stator retrieved previous cell type annotations, yet also found multiple states for each previous annotation (Fig. 5B). For example, cells previously annotated as being endothelial are uniquely enriched in 7 states (#4-6, #23, #32-34; green box in Fig. 5B). To cross-reference the same states in panels A and B we use an alluvial plot. Rather than calculating enrichments for disease status (panel A) or cell type annotations (panel B) separately, Stator also can perform an enrichment analysis for cells with Stator state labels with both previous cell type and disease/uninjured status annotations (panel C). This shows, for example, that whereas state #4 is enriched among cirrhotic sample cells (Fig. 5A) and among annotated endothelial cells (ECs) (Fig. 5B), it is enriched not just in cirrhotic but also uninjured ECs (Fig. 5C).

**Figure 5:**
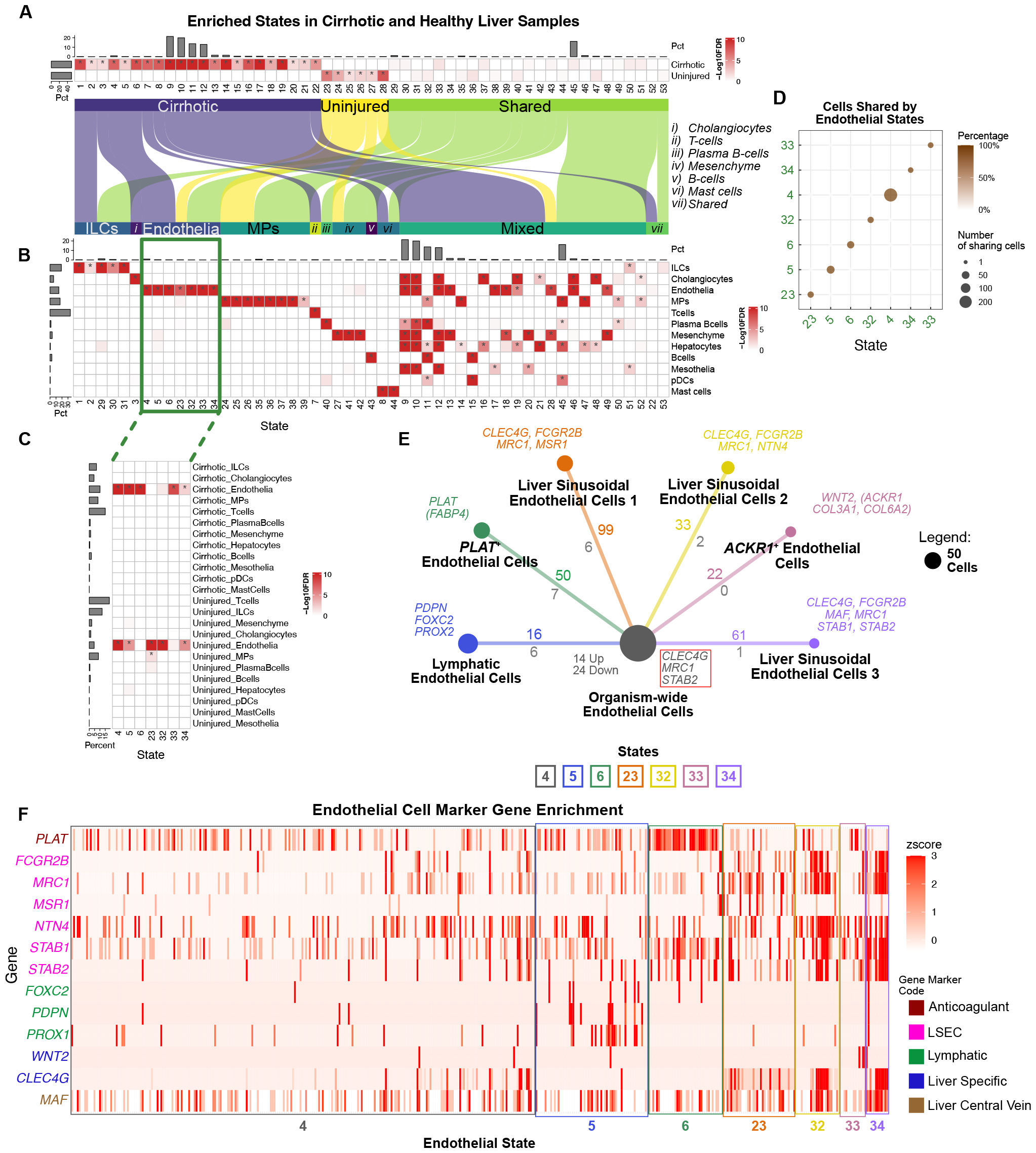
Stator states in cirrhotic and healthy human liver cells previously annotated by Ramachandran et al. (2019). (A) States (columns) enriched in single cells from cirrhotic or healthy liver samples (rows). (B) Heatmap showing states significantly enriched in these cells’ previous annotations (indicated by asterisks). Seven states (#4, #5, #6, #23, #32, #33, and #34) are significantly enriched only in the endothelial cell (EC) annotation (green box). (C) States significantly enriched in both cirrhotic/uninjured status and a previous cell type annotation (indicated by asterisks). (D) Virtually all cells with previous EC annotations are labelled with just one of the 7 EC specific cell states. These states were not detected by the original study (Ramachandran et al., 2019) or differential abundance analysis by Milo (Dann et al., 2022). (E) Numbers of cells labelled with EC states (#4, #5, #6, #23, #32, #33 and #34); areas of circles are proportional to their number (see panel legend). For states #5 to #34, numbers of significantly differentially expressed genes between cell cycle state pairs (*i.e*., s2s-DEGs) are indicated relative to state #4; colours refer to the state showing higher expression. Coloured numbers indicate significantly differentially expressed genes in cells labelled with state #4 compared to cells in any other EC state (*i.e*., s2o-DEGs); numbers of significantly differentially expressed genes between state #4 and all other EC states (increased and decreased expression) are shown in grey. Colour coded marker genes used to annotate cell states are provided adjacent to each state’s circle; a red box contains 3 genes whose expression is decreased in state #4 relative to the other EC states. (F) Heatmap of expression levels (z-scores) for marker genes used to annotate each EC state. Genes are grouped and colour coded by their associated annotation from the literature (Supplementary Table 2). The five categories of gene markers are colour coded as indicated in the panel legend. Cells (columns) are enclosed within a coloured box designating the EC state labelling that cell.

None of the 7 EC-labelled Stator states co-occur in 5 or more cells (Fig. 5D) suggesting that each of the 7 represents a distinctive EC subtype. Cross-referencing these states’ s2s-DEGs (Supplementary Table 21) to literature EC gene markers (Przysinda et al., 2020; Trimm and Red-Horse, 2023) predicted state #5 to be *PDPN, FOXC2*, and *PROX2* -expressing lymphatic-specific ECs; state #6 to be a subpopulation expressing *PLAT*, whose protein level is increased in patients with liver disease (Leiper et al., 1994); states #23, #32 and #34 to be liver-specific liver sinusoidal ECs (LSECs); and, state #33 to be a *WNT2, COL3A1, COL6A2* and *ACKR1* -expressing fibrotic niche subpopulation whose cirrhosis-associated expansion was discussed in (Ramachandran et al., 2019) (Fig. 5E). Using s2o-DEG analysis (Supplementary Table 22), the most populous state (#4) appears to label ECs that are not tissue or organ specific. Differential expression of these EC subtype marker genes across these ECs is illustrated in Fig. 5F. In summary, Stator has labelled cell subtypes from among a previously homogeneous set of ECs that were scarce in this dataset (*<*2.5%), demonstrating how rare subtypes can be discerned even within an under-represented cell type.

### 2.5. Stator recapitulates cancer cell types and NMF state annotations, yet at higher resolution

Finally, we applied Stator to a cancer data set, defining cancer cell states that were then compared against two sets of annotations defined previously by (i) clustering (Stuart et al., 2019) and comparison against reference datasets using SingleR (Aran et al., 2019), or (ii) non-smooth, non-negative matrix factorization (nsNMF; (Pascual-Montano et al., 2006)). For this analysis, 51 Stator states were predicted from 14,698 cells derived from 4 patients’ hepatocellular carcinoma (HCC) samples (Barkley et al., 2022) (Supplementary Table 23). These states were enriched for 11 of 12 cell types previously annotated using clustering and SingleR (Barkley et al., 2022) (Fig. 6A); the exception, epithelial cells, were low in number (*n* = 21). As before, Stator resolved single cell types into multiple subtypes, for example a single B-cell annotation into 12 sub-states. Myeloid lineage (macrophages, dendritic cells [DC] and neutrophils) states and lymphoid lineage (T cells, natural killer [NK] cells and B cells) states were distinct, highlighted in Fig. 6A by blue and red boxes respectively. Stator states were often easily annotated by their d-tuple genes. For example, state #43’s d-tuple genes contained *CD4* and other T cell markers; the myeloid lineage state #32 [*PLBD1* +, *SPI1* +, *LYZ* +, *MS4A6A*+] is in part defined by *MS4A6A*, a known marker for neutrophils, macrophage and dendritic cells (Franzén et al., 2019); and the lymphoid lineage state #48 [*IGHG4* +, *IGKC* +, *FGFBP2* +, *IGHG1* +] is largely defined by immunoglobulin genes, known markers for terminally differentiated B cells, *i*.*e*., plasma cells (MacParland et al., 2018).

**Figure 6:**
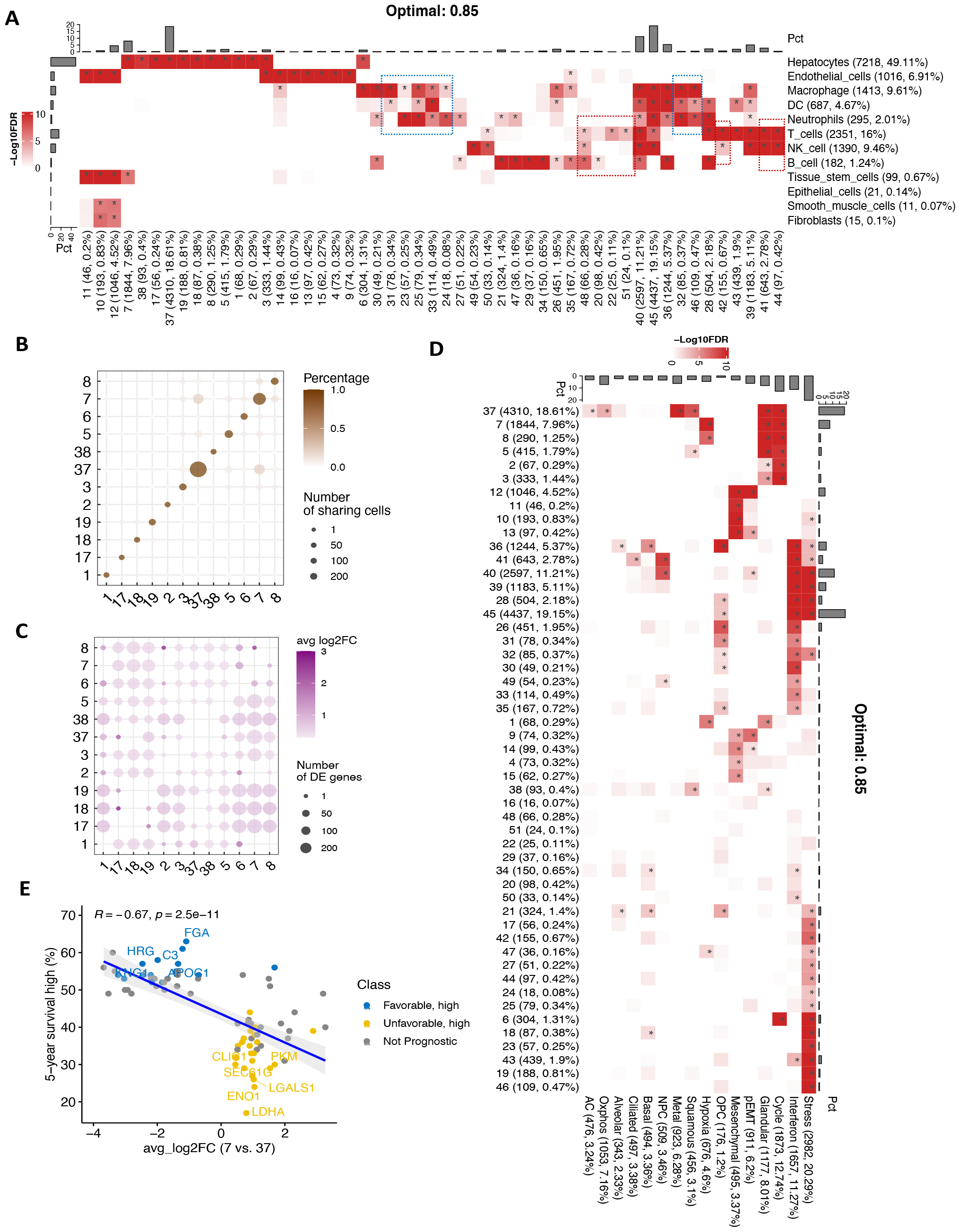
Stator identifies HCC cell types and states at higher resolution than clustering followed by SingleR (A) and NMF followed by expert annotation (D) respectively. **(A)** Heatmap showing significant enrichment (asterisks) among 51 Stator states with 12 cell type annotations previously defined by clustering followed by SingleR annotation (Barkley et al., 2022; Aran et al., 2019; Stuart et al., 2019). Stator identified multiple subpopulations for previously identified single cell types; for example, 11 Stator states occur unusually often only among cells previously annotated as hepatocytes. **(B)** Since Stator allows for cells to exist in multiple states, hepatocyte states can co-label single cells. Nevertheless, in the main these cells are only labelled with single Stator states. **(C)** Numbers of s2s-DEGs and their mean log2-fold change between Stator states enriched in cells previously annotated as hepatocytes. The 12 hepatocyte-enriched states are transcriptionally distinguishable. **(D)** Statistically significant enrichment (notified by asterisks) of Stator states (Y-axis) in cells previously annotated (Barkley et al., 2022; Gaujoux and Seoighe, 2010; Puram et al., 2017) into 16 NMF-defined states (X-axis). **(E)** Two Stator states are differentiated by genes that are predictive of liver cancer patient survival. Mean s2s-DEG expression fold-change (X-axis) for state #7 over #37 plotted against the percentage of 5-year survival (Y-axis) for TCGA patients whose expression of this gene lies above a pre-determined threshold (Uhlén et al., 2017).

The most populous state, #45, labels C1Q+ macrophages, an immunosuppressive population (Revel et al., 2022), annotated because 24 of 25 gene markers for these macrophages (cluster 10 of (Sharma et al., 2020)) are s2s-DEGs relative to state #40 (Supplementary Table 24). These are tissue-resident, rather than tumour-associated, C1Q+ macrophages because state #45 cells significantly more highly express *FOLR2*, rather than *TREM2*, relative to state #40 (Revel et al., 2022).

Twelve Stator states were enriched among cells labelled previously as hepatocytes (Barkley et al., 2022). These states labelled largely distinct sets of cells (Fig. 6B) that are transcriptionally distinguishable, as evidenced for example by large numbers of s2s-DEGs (Fig. 6C, D). A large minority (8%-23%) of these states’ s2s-DEGs are not expressed in normal hepatocytes (Methods), thereby reflecting their transformed status. The 12 transformed hepatocyte states showed considerable cell cycle gene expression heterogeneity. For example, State #7 expressed 6 cell cycle genes (*BIRC5, CCNA2, CCNB2, CDK1, TOP2A* and *UBE2C*) significantly more highly than the most populous State #37 (Supplementary Table 25). Other states (#17, 18, 19, 38) showed lower expression of these genes. These 6 cell cycle genes are rarely expressed in normal liver samples (Methods) and each gene’s high expression is known to be prognostic of worse outcome in liver cancer (Uhlén et al., 2017).

In the previously published analysis, these HCC cells were annotated both by type and state (Barkley et al., 2022). To investigate whether Stator could resolve these cells more finely, we analysed only those with both ‘Hepatocytes’ and ‘Cycle’ annotations, finding them to be enriched in 7 Stator states, most frequently in #37 (39% of 1,650 cells) and/or #7 (36%) (Fig. S14). Despite their previous identical annotation, cells in these 2 Stator states are transcriptionally divergent, with 78 s2s-DEGs separating them (Supplementary Table 26). State #37 more highly expressed transcripts that are abundant in normal hepatocytes (34 of 34 s2s-DEGs *e*.*g*., *AHSG, PLA2G2A, CYP2E1* and *HPD*) whereas state #7 more highly expressed genes that are rarely or never expressed in normal hepatocytes (13 of 44 s2s-DEGs, *e*.*g*., *TFF1, TFF2, TFF3* and *NDUFA4L2*). This suggests that Stator state #7 cells are in a more advanced state of cellular transformation than #37 cells.

To test this hypothesis we used TCGA liver cancer prognosis data (Uhlén et al., 2017, 2015), plotting s2s-DEGs’ mean expression fold-change (state #7 over #37, this study; Fig. 6E, X-axis) against the 5-year percentage survival rate (Y-axis) for TCGA patients whose expression of this gene is above a pre-determined threshold (Y-axis, (Uhlén et al., 2017)). This showed that genes that are more highly expressed in state #7 over #37 tend to be those genes that are more highly expressed, at diagnosis, in liver cancer samples of patients with lower survival rates. Conversely, genes that are more highly expressed in state #37 over #7 tend to be genes that are more highly expressed in liver samples of patients with higher survival rates. In summary, Stator has revealed previously unappreciated HCC cancer states whose differential expression involves genes that are predictive of patient survival.

## 3. Discussion

In flow or mass cytometry, cells are typed from heterogeneous samples by the presence or absence of one or more labelled antibodies, all chosen a priori. Single cell transcriptomics types cells in a more data-driven manner, first clustering cells’ transcriptomes and then identifying genes that are differentially expressed between clusters. Nevertheless, this approach: (i) types cells using only gene expression and not non-expression, (ii) when combined with projection to a low-dimensional 2D space for visualisation, suffers from significant distortions (Chari and Pachter, 2023), and (iii) relies on localisation of cells in expression space. The latter presents an additional problem because cells need then to be given multiple labels, for example type, sub-type, cell cycle phase, maturity and activity (Kotliar et al., 2019), which, however, need not be localised in expression space.

We have shown how Stator provides multiple type and state labels for heterogeneous samples of single cells by identifying combinations of genes that are co-ordinately expressed and/or nonexpressed in single cells. These states are provably different from correlations Fig. S1. Rather than supplying a single cell type marker, which can mislead as when *Aldh1l1* or *Gfap* only is used for astrocyte identification (Akdemir et al., 2020), and rather than requiring a hierarchy of cell types (Zeisel et al., 2018), Stator differentiates cells by primary (cell type), secondary (sub-type) and tertiary (cell state, activity, cell cycle phase, or maturity) markers. States are treated as entities drawn from a continuum, which more accurately reflects the progression from radial precursor cells to neurons, for example, due to processes of activation and differentiation (Dulken et al., 2017; Yuzwa et al., 2017).

Gao et al. (2022) recently solved the issue of selective inference bias, or *double-dipping*, specifically when cells are clustered by optimising their transcriptional differences before calculating their transcriptional differences. Each of these two operations occurs on gene expression space. Stator clusters not cells, but rather d-tuple gene signatures, prior to s2s-DEG analysis. Even if present, Stator will mitigate selective inference, at least in part, by differences not being maximised on the same space, and by demanding significant s2s-DEGs to not just be d-tuple genes, when states are declared to be transcriptomically non-identical. The Gao et al. method is also not immediately applicable here due to its reliance on clustering algorithms that compute Euclidean distances, whereas Stator relies on Dice similarity.

Due to current computational constraints, Stator is limited to approximately 1,000 HVG and 40,000 cells to estimate higher-order *n*-point interactions (*n* = 2, 3, …, 7). Estimation of conditional dependencies contributes most to computational cost, so Stator’s efficiency and accuracy could be greatly improved as new causal discovery methods are developed. In addition, accuracy could be improved by integrating biological knowledge into the dependency graphs. The limitation of up to 7-point interactions is statistical rather than computational: we did not find evidence for significant 7-point interactions in the datasets analysed. Stator takes advantage of sparse gene-by-cell matrices, and so is not intended for analysing deep coverage transcriptomes until more sophisticated binarisation schemes are explored (e.g., (Li and Quon, 2019)). Other challenges relate to how Stator predictions should be interpreted, particularly those states lying on a continuum whose biology is poorly understood. Further, the resolution (*i*.*e*., Dice similarity) at which states should be defined and can be interpreted will vary by data set. Lastly, conditioning on absent gene expression in the Markov blanket (Eq. 1, Methods) may overlook some states despite large numbers of biologically plausible states being returned.

Stator results show that a wealth of biological information can be inferred from the higher-order statistics of single-cell expression data. Evidence exists for higher-order and combinatorial genetic interactions (Kuzmin et al., 2018; Antebi et al., 2017; Arnosti et al., 1996; Watkinson et al., 2009) and pairwise quantities at different pseudotimes have been investigated (Ghazanfar et al., 2020). Nevertheless, the biological value of higher-order statistics in single-cell gene expression has not previously been shown.

The picture emerging from applying Stator is of cells adopting a spectrum of states (or colours, in this metaphor) with their primary colour representing their strongest transcriptomic signature, most often indicating cell type. Differential expression between cells of the same type, but in two different states, filters out their primary colour thereby revealing secondary colours, representing cellular dynamics differences. This metaphor can be continued with respect to tertiary and quaternary colours, representing even more finely resolved aspects of cell state.

Stator can be applied to a variety of scRNA-seq data sets in biomedicine, including those with a temporal label (*e*.*g*., developmental or disease progression), as well as data from different individuals, to compare and contrast cell states of individuals with different disease progression trajectories, or responders and non-responders to therapy. Finally, Stator’s general methodology can also be applied to other datasets with variables that are binary or can be approximated well by binarisation, such as disease comorbidities, scATAC-seq, or sparse single cell proteomics.

## 4. Methods

### 4.1. n-point interaction estimation

In previous work we developed a model-independent estimator of higher-order interactions amongst binary variables (Beentjes and Khamseh, 2020). Here, we refer to the multiplicative interaction in (Beentjes and Khamseh, 2020) as Model-Free Interaction (MFI) due to its definition being without reference to any subjective parametric model, but in terms of probabilities and their expectation values. Similar notions (for 2-point interactions) have been proposed in the statistics literature (Hernan and Robins, 2023; VanderWeele and Knol, 2014). For completeness, we summarise the main definitions and interpretations of MFIs (Beentjes and Khamseh, 2020) below. A 2-point MFI is defined, and can be rewritten, as follows:

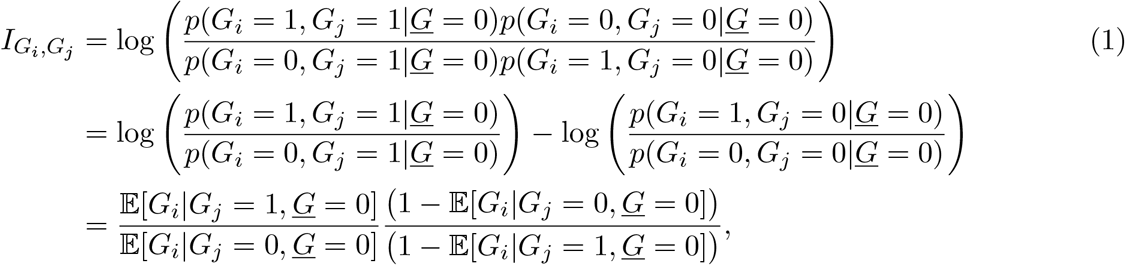

where <u>*G*</u> is the set of all other genes, aside from *G*_*i*_ and *G*_*j*_, that are not independent of *G*_*i*_ and *G*_*j*_. The first line in Eq. 1 has the interpretation of a generalised conditional log-odds ratio and is symmetric in *G*_*i*_ and *G*_*j*_. The second line provides the following interpretation: “Does the likelihood of gene *G*_*i*_’s expression being *on* vs *off* depend on the status of gene *G*_*j*_’s expression?”. To elaborate further, the first term represents the likelihood of gene *G*_*i*_ being *on* vs *off*, whilst gene *G*_*j*_ is *on*, while the second term represented the same quantity with gene *G*_*j*_ is *off*. If the expression of the two genes *G*_*i*_ and *G*_*j*_ are completely independent of each other, then these two terms cancel and result in a zero interaction as desired. The third line represents the same quantity in terms of expectation values, which are then taken as averages over the data for estimating the interactions. Uncertainties in these estimates are quantified via the bootstrap procedure (Efron, 1979). In (Beentjes and Khamseh, 2020) we generalised this definition and estimator to *n*-point interactions. For example, a 3-point interaction, where *p*(*G*_*i*,*j*,*k*_ = 1, 1, 1) is shorthand for *p*(*G*_*i*_ = 1, *G*_*j*_ = 1, *G*_*k*_ = 1|*<u>G</u>* = 0) and so on, is defined as follows:

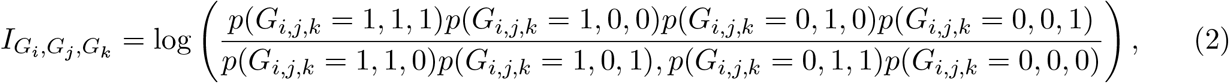

and has the interpretation of whether the expression status of a third gene, *G*_*k*_, changes the 2- point interaction between *G*_*i*_ and *G*_*j*_ expression. We presented previously (Beentjes and Khamseh, 2020) that this definition recovers, in a data-driven manner, known ground truth interactions in statistical physics systems such as the Ising model, and more generally energy-based models, as well as any other Markovian complex system. We further demonstrated that our MFI definition, used to directly estimate the interaction, results in the same estimate as when training a Restricted Boltzmann Machine, both analytically and numerically within statistics. The advantage of the MFI direct estimation on binary data is its model-independent definition interpretability and its avoidance of having to fit the joint probability distribution amongst the variables. The latter is a much more complex quantity to estimate robustly than the combination of expectation values in the MFI estimator. Finally, we note that conditioning on <u>*G*</u> = 0 in Eq. 1 is equivalent to finding the ‘pure’ 2-point interaction between *G*_*i*_ and *G*_*j*_ without the influence of the other genes’ expression. Note that <u>*G*</u> need not contain the set of all other genes when estimating the interaction. Indeed, it is sufficient to only condition on the Markov blanket (MB) of *G*_*i*_ and *G*_*j*_, *i*.*e*., the smallest set of genes <u>*G*</u> conditional on which *G*_*i*_ and *G*_*j*_ are independent of all other genes. Once conditioned on the MB, the information from other genes no longer influences the interaction between *G*_*i*_ and *G*_*j*_. Therefore, restricting <u>*G*</u> to only contain the MB of genes for each pair *G*_*i*_ and *G*_*j*_, improves statistical power, whilst simultaneously ensures that the 2-point interaction remains stable by measuring the direct dependence between *G*_*i*_ and *G*_*j*_, rather than indirect correlations. The same argument holds for higher-order interactions. Fig. S1 presents a comparison between MFIs, correlation, partial correlation and mututal information, computed on data generated from a set of DAGs in accordance to Fig. S2, first presented in (Jansma, 2023a). The set of MFIs is distinct between distinct DAGs, whereas other dependence metrics are only able to distinguish some, but not all, distinct DAGs.

Currently, performing conditional independence tests amongst all groups of genes to determine their MBs, is statistically and computationally prohibitive. For this reason, we restrict the estimation of *n*-point interactions to the top 1,000 HVGs, after quality control. Stator then infers the MBs of the HVGs via a hybrid Bayesian network inference technique (Kuipers et al., 2022) which sequentially performs (conditional) independence testing, starting from a fully connected undirected graph of genes (Peter-Clark algorithm (Spirtes et al., 2001)), followed by a score and search MCMC approach to obtain the optimal completed partially directed acyclic graph (CPDAG), introduced in Kuipers et al. (2022). We emphasise that we do not claim any causal inference or regulatory relationships amongst these genes based on the inferred network. Instead, we utilise this algorithm to infer a gene signature dependence network structure to obtain the MB and estimate higher-order interactions with sufficient statistical power, with the final aim of inferring cell (sub)types and states. Inferring this dependence network massively reduces the search space for potentially significant interactions. For run-time considerations see Supplementary Material 7.1.

Finally, we note that MFIs are symmetric. Therefore, when estimating, *e*.*g*., a 2-point interaction using line 3 in Eq. 1, one can choose to estimate the terms 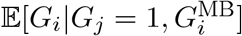 or 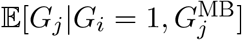, whichever results in the greatest statistical power, *i*.*e*., when either the MB of *G*_*i*_ or *G*_*j*_ is smaller, or more generally, when the MB of *G*_*i*_ or *G*_*j*_ is more populated.

Having identified the set of MBs, Stator then estimates up to 7-point interactions amongst the genes in the expression data. The 2-point interactions are estimated between all pairs of genes, the 3-, 4-, and 5-point interactions are estimated amongst all gene tuples that are in each other’s MB (the interaction amongst Markov disconnected genes vanishes (Jansma, 2023a)), and 6- and 7-point interactions are calculated amongst genes that are in the MB of a tuple of genes with a significant 5- or 6-point interaction. Every interaction is estimated using the smallest possible MB.

In order to prioritise candidate interactions for the next step (“Deviating gene tuples (d-tuples)”), each interaction is estimated 1,000 times by bootstrap resampling the data. An interaction is prioritised as a ‘non-zero’ candidate for the next step if the fraction *λ* of bootstrap estimated interactions with a different sign from the original estimate is less than 0.05. This procedure is more permissive than testing for the hypothesis that the 95% two-sided percentile bootstrap confidence interval does not contain zero. For the data sets studied in this work, we verify numerically that this procedure is equivalent to demanding 90 − 95% confidence, depending on the order of the interaction.

### 4.2. Deviating gene tuples (d-tuples)

In a finite sample of *N* cells, the observed frequency Φ_*s*_ of a tuple *s* = *{s*_1_, …, *s*_*n*_*}* of *n* independently expressed binarised genes is binomially distributed as:

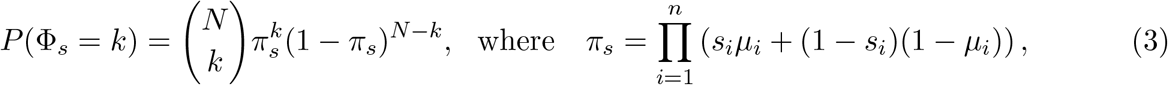

and *µ*_*i*_ is the mean expression of gene *i* across all cells under consideration (*i*.*e*., the cells for which the relevant MB is zero). Equation 3 describes the null hypothesis that the observed cell counts are the result of independently expressed genes, and gives the expected number of cells under this null: 𝔼[Φ_*s*_] = *π*_*s*_*N* . An observation Φ_*s*_ = *ϕ*_*s*_ of one of the 2^*n*^ joint states of *n* genes can be assigned a p-value:

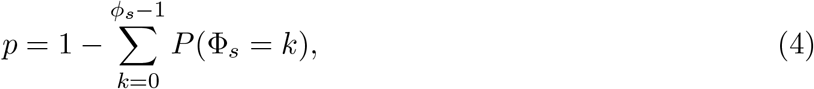

and log 2-fold change, or deviation:

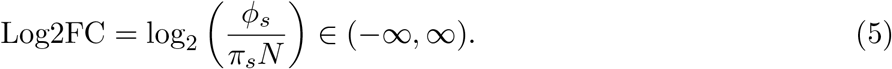

The p-values are calculated for all tuples with a positive Log2FC, and corrected for multiple hypothesis testing with the Benjamini-Yekutieli procedure (Benjamini and Yekutieli, 2001). A non-zero interaction can thus have one or more *deviating-tuples (d-tuples)*, those tuples of genes that significantly deviate from the null hypothesis. Since a non-zero interaction reflects a higher-order dependency in the data, its d-tuple describes the gene expression patterns that are (at least in part) responsible for this dependency. The set of cells that have the *n* genes in that particular expression state—ignoring the state of the MB—form the associated set of cells. Note that cells carrying a certain combination of d-tuples need not cluster in expression space: whilst these cells all share a particular gene expression pattern among the *n* genes, the expression of all other genes can vary greatly. This makes it in principle possible for a cell state to be widely dispersed in expression space.

### 4.3. Hierarchical clustering of d-tuples

Given all d-tuples, Stator creates a cell-by-d-tuple matrix, with binary entries 1 or 0, representing whether or not a cell contains a particular gene d-tuple. Stator then hierarchically clusters these d-tuples (rather than cells) based on a notion of distance, here the Sørensen–Dice coefficient, to identify d-tuples that more commonly co-label the same cells. This hierarchy of separation among d-tuples can be visualised in a dendrogram. Note that the Sørensen–Dice coefficient, sometimes referred to as the Dice similarity coefficient, is not a distance metric because it does not satisfy the triangle inequality. More specifically, the Dice similarity between two boolean vectors *X* and *Y* is defined as:

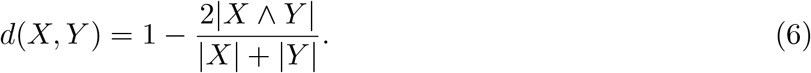

In order to group the cells together, we cut the dendrogram at a Dice similarity that, by default, is set at the value that maximises the weighted modularity score of the resulting clustering (Newman, 2006), where a pair of d-tuples is assigned an edge weight of one minus their Dice similarity. At the set Dice similarity threshold, cells expressing these gene d-tuples are grouped together forming Stator states. In particular, cells can exist in multiple multiple Stator states depending on different gene signature similarities. Lowering the Dice value threshold increases granularity, the resolution by which states are predicted, which we have shown, in some instances (*e*.*g*., Fig. S12), to better resolve subtypes or substates for large and transcriptionally heterogeneous groups of cells.

### 4.4. Stator pipeline

Stator is a Nextflow pipeline (written using Nextflow version 21.04) that consists of a main Nextflow script (DSL1) managing a number of Python and R scripts and modules (see Figure S4 for an overview of the pipeline). Stator aims to balance modularity and ease-of-use with flexibility, so is fully containerised (Docker images hosted on Dockerhub) and allows the user to specify different preferences and settings in a separate json file, meaning that it should run reproducibly on any Sun Grid Engine compatible cluster. The only file that has to be supplied by the user is a .csv file (called `rawDataPath` in the json settings file) containing the expression data of <u>*G*</u> genes (columns) and *C* cells (rows), where the first row contains the column/- gene names. Optionally, the user can provide a file userGenes that contains the names of genes that should be included in the final analysis regardless of their variability, a file `genesToOne` containing genes whose Markov blanket state should be 1 instead of 0 (not used in this paper) which allows for conditioning on different Markov blanket states, and a file doubletFile containing a Boolean exclusion list, for example based on a doublet annotation, that indicates which cells should be excluded, regardless of other QC metrics. The user should further specify the total number of cells (`nCells`) and genes (`nGenes`) to be used in the analysis.

The pipeline then initiates the first process in the pipeline, defined in the makeTraining- Data.py script. By default, Stator assumes that the data is already quality controlled (QCed) and only performs very basic data preparation (specified by the setting `datatype=‘agnostic’`): all cells specified by doubletFile are excluded, PCA and UMAP embeddings are calculated, and up to `nGenes` genes are included, starting with those specified in `userGenes`. A total of nCells is then randomly selected for downstream analysis. Alternatively, Stator can run in `datatype=‘expression’` mode and perform basic scRNA-seq QC, where parameters such as the threshold of mitchondrial gene reads can be set by the user. `In expression` mode, Stator first includes the userGenes, but then adds the most highly variable genes until nGenes are included. The final count matrix of size `nCells` *×* `nGenes` is then binarised and sent to the next process.

Stator then aims to estimate the graph of conditional dependencies among the genes. It does this by generating a first guess using a parallelised implementation of the Peter-Clark (PC) algorithm (parallelPCscript.R, based on (Le et al., 2016)). The PC-algorithm starts with the fully connected graph of dependencies, and then iteratively performs dependency tests among connected pairs, removing an edge when no evidence for dependence is found (delaying removal until all tests are done to ensure order-independence. In addition, we use the majority-rule suggested by (Colombo et al., 2014)). Somewhat counter-intuitively, a larger significance threshold for the dependency tests corresponds to a more conservative estimate, since preserving more edges will result in larger MBs which are necessarily more conservative. The default threshold is set at *p <* 0.05, not corrected for multiple hypothesis testing, but can be adjusted by the user. Reducing this threshold makes the estimate less conservative, but can significantly speed up the estimation procedure by eliminating more edges and reducing the size of the estimated MBs. This initial guess is then iteratively improved upon using the score-based MCMC method outlined in (Kuipers et al., 2022) (iterMCMCscript.R). This method is based on an efficient exploration and scoring of the space of possible DAGs, and allows new edges to be introduced into the initial guess if they significantly increase the score. The CPDAG equivalence class corresponding to the graph found by parallelPCscript.R is used as the starting point, and the script iterates until increasing the search space no longer increases the score. To be as conservative as possible in our estimates, the final MBs used in all downstream analysis are those based on the full final search space on which this algorithm terminated (not, for example, only the *maximum-a-posteriori* estimate or its associated CPDAG).

Using these MBs, all 2-to-5-point interactions are calculated among genes that are mutually Markov connected (`calcHOIsWIthinMB.py`). By default, uncertainty is quantified by bootstrap resampling, but this can be done more efficiently using an estimate for the asymptotic error rate of the MFIs by setting asympBool=1 in the settings; agreement with bootstrapped confidence intervals was confirmed previously (Jansma, 2023b).

The higher-order interactions are analysed (`createHOIsummaries.py`) and used to calculate the significant d-tuples and final Stator states (`identifyStates.py`). In addition, if there are interacting 5-tuples that are Markov connected to additional genes, a targeted search for 6- and 7-point interactions is performed. Run time using reasonable settings is discussed in Appendix 7.1.

Stator’s output includes files containing both the binarised and unbinarised QCed expression data, a list of all d-tuples, and cell embedding coordinates. These can then be used for further downstream analysis, for which we provide an R Shiny app. More information on the various settings available to the user, as well as a complete list of output files, is available at https://github.com/AJnsm/NF_TL_pipeline/tree/main.

### 4.5. Stator’s R Shiny App

The Stator App was implemented as a web application for downstream analyses, using the R Shiny package (v1.7.4) from R studio (shiny.rstudio.com). As an open-source application, the code is available through GitHub at github.com/YuelinYao/MFIs. The Docker container image could be found from Dockerhub: hub.docker.com/r/yuelinyao120/statorapp. Stator App is hosted at shiny.igc.ed.ac.uk/MFIs/. A complete list of packages used can be found at github.com/YuelinYao/MFIs/blob/main/renv.lock. The app consists of twelve main panels (*About, Table, Heatmap-Cells, Heatmap-Genes, GO & KEGG, Using rrvgo, Upset Plot, DE analysis, Find Markers, Automatic Annotation, Markov Blanket, UMAP Plot*).

#### 4.5.1 Data upload and file input

The app begins with an About page, providing general information about the app and a tutorial on its use. It includes a liver cancer dataset (Barkley et al., 2022) already uploaded, and users can upload their own files in ‘.csv’ format (size *<*100GB running from server); most of these are output files from the Stator Nextflow pipeline. The Tutorial explains how to prepare a dataset, and provides information on the app’s parameters and statistical tools.

#### 4.5.2. Summary Table

The app generates a statistics summary table by filtering and clustering significantly deviating tuples (d-tuples) after file upload and parameter setting. The minimum enrichment factors in Log2 transformation has a default value of 3, *i.e*., 8-fold change, the minimum number of cells labelled by each d-tuple has a default value of 0, and the FDR is by default set to 0.05. These parameters are used to filter d-tuples. The Dice similarity is employed for hierarchical clustering of d-tuples. This table presents tuple genes and their state, along with their respective enrichment factor in log2 transformation, adjusted enrichment p-value, the number of cells labelled by each d-tuple, and the cluster that includes this d-tuple. The d-tuples in each cluster define a Stator cell state.

#### 4.5.3. Cell states with external annotations

The Shiny App offers the ability to explore cells and genes in each state using externally provided annotation through Heatmap-Cells and Heatmap- Genes panels, respectively. Users can additionally specify the type of analysis they wish to perform, such as annotation term enrichment analysis (over-representation test), depletion analysis (underrepresentation test), or a two-sided Fisher’s exact test.

a. Enrichment analysis for cells: Enrichment analysis allows users to test for the enrichment of external annotation terms in Stator states. We use the following notation: *N* : Total number of cells. *m*: Number of cells (of total *N*) in a given Stator cell state. *k*: Number of cells (of total *N*) in the external annotation. *q*: Number of cells shared between a given Stator cell state and an external annotation. The corresponding random variable is denoted by *X*. The null hypothesis is that the observed overlap between the identified cell state and the external annotation is no greater than is expected by chance. The *p*-value is calculated as the probability of observing more overlapping cells than expected under this null hypothesis: 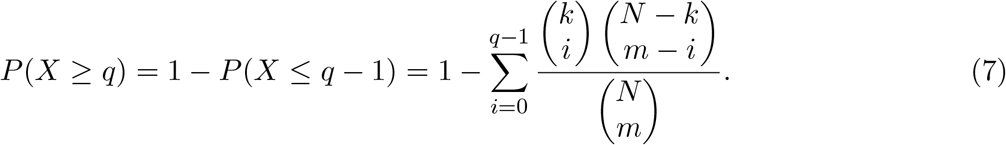 The probability is computed with the R function: `phyper(q-1, m, N-m, k, lower.tail = FALSE, log.p = FALSE)` Once the *p*-value is computed for all pairs, we use the Benjamini and Hochberg (BH) method (Benjamini and Hochberg, 1995) for correcting for false positives arising from multiple tests. The corrected *p*-values are transformed by taking the negative logarithm (base 10) before then being visualised as a heatmap, using ComplexHeatmap (v2.14.0) (Gu et al., 2016; Gu, 2022).
b. Depletion analysis for cells: the null hypothesis is that the observed overlap between the identified cell state and the external annotation is no fewer than would be expected by chance. The *p*-value is computed with the R function: `phyper(q, m, N-m, k, lower.tail = TRUE, log.p = FALSE).`
c. The two-sided Fisher’s exact test: As an option, a two-sided Fisher’s test may be performed with following R function and the heatmap provided from this test is coloured by the log10 transformed odds ratio: `fisher.test(matrix(c(q, m-q, k-q, N-m-k+q), 2, 2), alternative=‘two.sided’)` A similar statistical test is performed, using the Heatmap-Genes function, to test for the overlap between an externally supplied gene list with genes listed among Stator state d-tuples.

#### 4.5.4. Gene Ontology and KEGG Pathway Enrichment analysis

The Shiny App allows users to perform Gene Ontology (GO) and KEGG Pathway Enrichment analysis for d-tuple genes in each Stator state using the R package clusterProfiler (v4.6.2) (Yu et al., 2012). Users can specify the cell state(s) of interest and reference genome for the dataset (e.g., `hsapiens_gene_ensembl, org.Hs.eg.db`, and hsa for human), or background genes. Significantly enriched terms (FDR *<* 0.05) are displayed in the app. The app also implements Rrvgo (Sayols, 2023) to reduce redundancy of GO terms and for their visualisation as word cloud, treemap or scatter plots.

#### 4.5.5. Cell Upset plot

The app shows Upset plots, with rows corresponding to numbers of cells labelled by each state, and columns providing the number of cells labelled in common. This uses the ComplexHeatmap package (v2.14.0) (Gu et al., 2016) in R.

#### 4.5.6. Differential expression analysis

The Shiny App allows users to perform differential analysis: i) between two cell states, disregarding all cells co-labelled with both states (termed s2s) from the DE analysis tab, or ii) between cells labelled with a state and all cells without this label (termed s2o) from the Find Markers tab. Differential gene expression analysis was implemented using the FindMarkers function from Seurat (v4.3.0) (Stuart et al., 2019). Users can define log2 fold change and adjusted p-value thresholds. The app then displays an expression heatmap of differentially expressed genes, a volcano plot, a summary statistics table for differential expression, and Gene Ontology and KEGG term enrichment significance results for differentially expressed genes.

In the automatic annotation tab, users can provided a table of genes of interest, and the app will identify s2oDEGs for all Stator states, and automatically return the DEGs in the provided gene list for easy anotatation.

#### 4.5.7. Markov Blanket, MB

The app provides the functionality for users to extract and visualise the MB for a particular gene. For this visualisation it imports the inferred MCMC graph and extracts the MB covering all parents, children and spouses of this gene. This was implemented by the R package, igraph (v1.4.1)(Csardi et al., 2006).

#### 4.5.8. UMAP plot

The app allows users to visualise a cell state of interest within an uploaded set of UMAP cell coordinates. This was implemented using the DimPlot function from Seurat (v4.3.0) (Stuart et al., 2019).

### 4.6. Datasets

To showcase Stator’s prediction of cell types, subtypes and/or states in diverse normal and disease samples, we chose three diverse datasets: (i) Normal brain tissue E18 mice from the 10X Genomics ‘1.3 Million Brain Cells from E18 Mice’ dataset (10x Genomics, 2017), downloaded from https://www.10xgenomics.com/resources/datasets, (ii) human liver tissue from control and disease (cirrhosis) samples (Ramachandran et al., 2019), and (iii) human liver cancer (hepatocellular carcinoma) tissue from Barkley et al. (2022). Dataset (i) contains an unannotated Louvain clustering (60 clusters in total) (Blondel et al., 2008) of 1,306,127 transcriptomes distributed over 133 libraries, sequenced on an Illumina HiSeq 4000 using paired-end sequencing at a moderate read depth of 18,500 reads per cell, keeping only uniquely mapped reads. To annotate these clusters by cell type, we identified upregulated (with respect to all other cells) marker genes using the `R-function scran::findMarkers` (Lun et al., 2016). Cluster 7 had top 10 marker genes (all at FDR*<* 10^−10^) *{****Syt6***, *Gm27032, Slain1*, ***Pbx3***, *Rgs8, Fgf3, Nkx2-3, Otor*, ***Six3, Myh7*** *}*. Gene symbols shown in bold are listed on mousebrain.org/adolescent/genes.html (Zeisel et al., 2018) as markers for CNS-neurons, while the other genes are not markers for any cell type (except for *Rgs8* which marks trilaminar cells). Furthermore, when inferring markers against specific other clusters, *Dlx2, Dlx5* and *Dlx6os1* appeared as top markers; these genes control GABAergic neuron differentiation in developing mice (Petryniak et al., 2007). Cluster 10 had top 10 marker genes (all at FDR*<* 10^−6^) *{****Gm11627***, *Abhd4, Mpv17, Cldn10, Dhrs1, Thbs3*, ***Aldoc, Prdx6***, *Gm20515, Chil1 }*; gene names in bold show upregulated expression in radial glial cell precursors at E17.5 (Yuzwa et al., 2017). Although, radial glial cell precursors and astrocytes are challenging to distinguish by differential gene expression (Dulken et al., 2017), mature astrocytes are not abundant at this early developmental stage (E18) (Akdemir et al., 2020). We therefore concluded that clusters 7 and 10 are composed of neurons and radial glial cell precursors (RPs), respectively. To create a merged dataset containing both neurons and RPs, we first merged both clusters, and then downsampled these to 19,000 cells, of which 13,905 were neurons, and 5,395 were RPs.

Dataset (ii) was generated downsampled from 58,358 to 20,000 cells, specifically by sampling 10,000 cells from uninjured samples and 10,000 cells from cirrhotic samples. When Stator states were compared with expert annotations, lineage annotations from the original publication were used (Ramachandran et al., 2019). No cells annotated as “cycling” by Ramachandran et al. (2019) remained after sub-sampling. Stator states for dataset (ii) used a Dice similarity of 0.97, with a minimum 8-fold enrichment of tuples over expected, a maximum FDR corrected enrichment significance of 0.05, and a minimum of 10 cells labelled by each d-tuple.

Dataset (iii) was generated from a pan-cancer dataset by selecting the liver tumor type, resulting in 14,698 cells (Barkley et al., 2022). Three types of annotations were defined in the original study: a.) cell type by clustering (Stuart et al., 2019) and SingleR (Aran et al., 2019), b.) cell state by nsNMF (Puram et al., 2017; Gaujoux and Seoighe, 2010), and c.) malignant or not by inferCNV (Patel et al., 2014). We defined a gene as being normally expressed in untransformed hepatocytes when it was expressed (≥ 1 read) in *>* 0.1% of hepatocytes (Andrews et al., 2022). Stator states for dataset (iii) used a Dice similarity of 0.85, with a minimum 8-fold enrichment of tuples over expected, a maximum FDR corrected enrichment significance of 0.05, and a minimum of 0 cells labelled by each d-tuple.

### 4.7. Quality Control (QC) and expression binarisation

Data used as input to Stator was pre-processed using standard Quality Control (QC) best practice (Luecken and Theis, 2019). When doublet removal was not performed in a study, or this information was absent, we removed doublets using Scrublet (Wolock et al., 2019). We restricted analysis to the 1,000 most highly variable genes (HVG), quantified using Scanpy (Wolf et al., 2018), followed by binarisation of gene expression. Droplet-based protocols commonly result in sparse data with many dropouts. Justification for gene expression binarisation has been previously demonstrated for a variety of scRNA-seq analyses including dimensionality reduction, clustering, differential gene expression and pseudotime analyses (Bouland et al., 2023, 2021; Qiu, 2020). Following the literature, we binarise expression values, with genes without expression evidence as zeros, and those with evidence of expression as ones.

### 4.8. Comparison with clustering

We applied hierarchical clustering on the two mouse brain datasets to compare Stator with the conventional clustering approach. We processed and selected the 2000 most HVG to compute principal components (Stuart et al., 2019), and then the top 20 PCs were used to calculate the Euclidean distances. Specifically, we used Ward’s method for hierarchical clustering, which is based on minimizing the loss of information from joining two groups (Murtagh and Legendre, 2014).

Using the same data to both cluster cells and test the differential expression will result in an extremely inflated type I error rate (Gao et al., 2022). To compare Stator with robust clustering results, we applied a selective inference approach to test for a pairwise significant difference between two clusters (Gao et al., 2022). This approach protects against selective inference by correcting for the hypothesis selection procedure (Gao et al., 2022). We applied Bonferroni method to correct the p-values for multiple comparisons. Ideal clustering should result in a significant p-value for any pair of clusters. To declare the final number of distinct clusters, we take the largest number of clusters such that all clusters are pairwise significantly distinct as the total number of clusters is tuned.

## 5. Acknowledgements

CPP and JCW were funded by the MRC (`MC_UU_00007/15`). AK was supported by the XDF Programme from the University of Edinburgh and Medical Research Council (`MC_UU_00009/2`) and is supported by a Langmuir Talent Development Fellowship from the Institute of Genetics and Cancer, and a philanthropic donation from Hugh and Josseline Langmuir. AJ was supported by an MRC Precision Medicine Grant (MR/N013166/1). AJ thanks Øyvind Almelid for many helpful discussions on Nextflow. YY thanks Xinyi Jiang for improving the Dockerfile of the shiny app.

## 6. Competing Interests

No competing interests declared.

## 7. Supplementary Material

**Figure S1:**
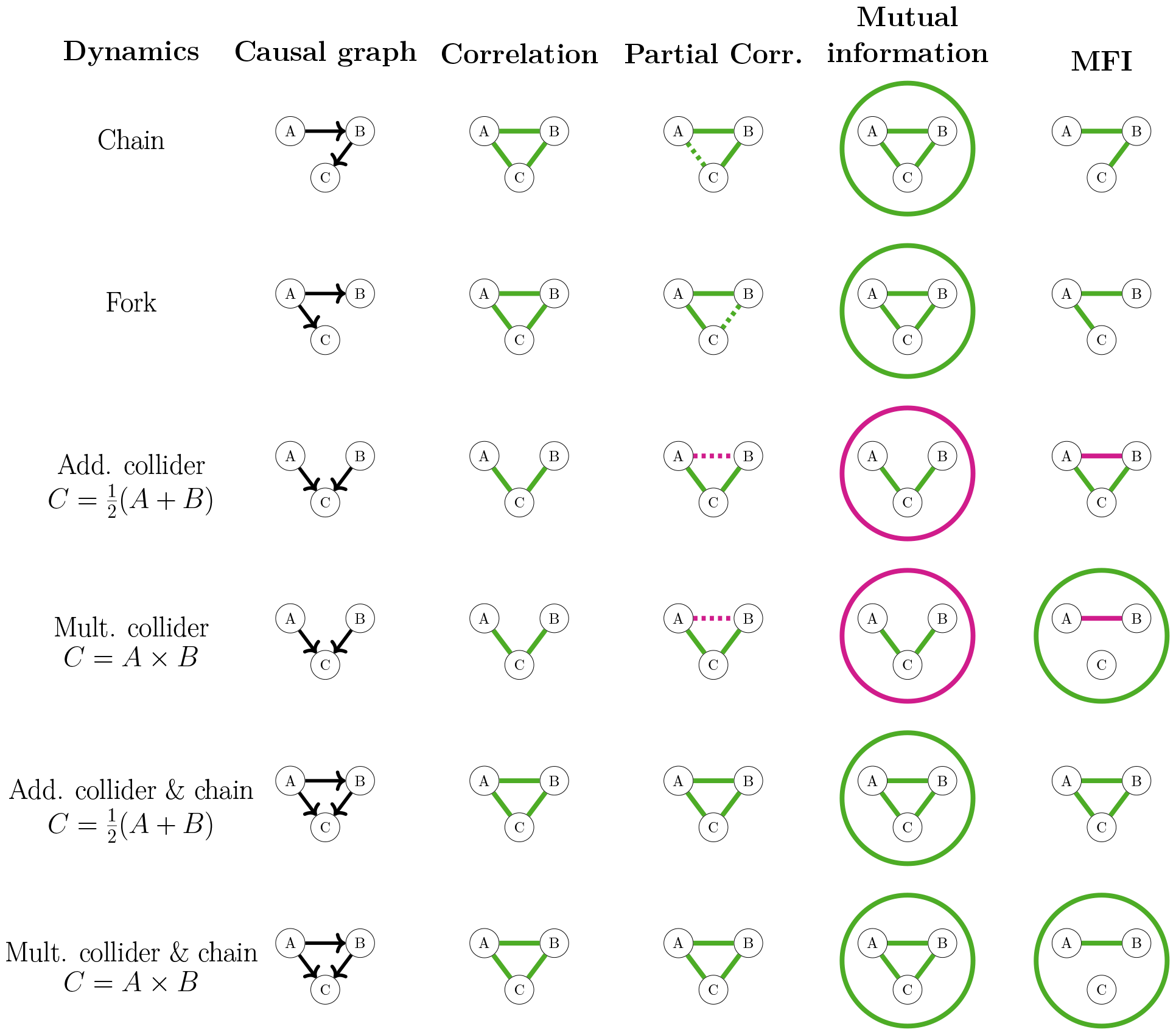
Extended Fig. 1. Comparison of MFIs with other estimators of dependence. Different causal dynamics lead to different association metrics, and only MFIs can distinguish all 6 scenarios and reveal the combinatorial effect of a multiplicative interaction. Green edges denote positive values, red edges denote negative values, circles denote a 3-point quantity, and dashed lines show edges that show marginal significance that depends on the level of simulated noise. Correlations and mutual information cannot distinguish between most dynamics, and while partial correlation can, for certain noise levels, identify the correct pairwise relationships, it falls short of distinguishing additive from multiplicative dynamics. See Fig.S2 for the simulation parameters and precise values. Reproduced from Jansma (2023a) with permission from the author.

**Figure S2:**
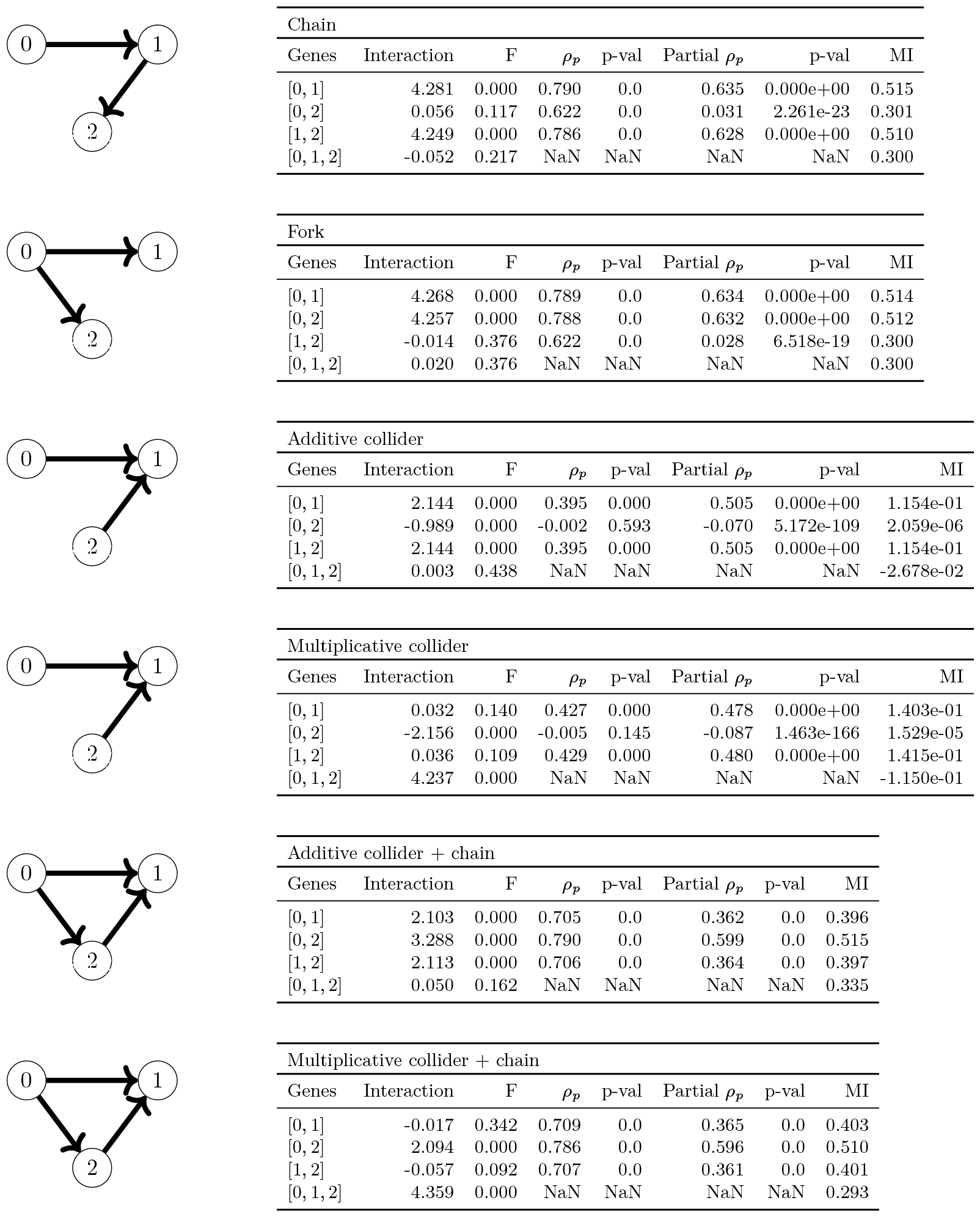
Here listed are the precise values that led to Fig. S1. From each graph, we generated 100k samples from a Bernoulli distribution with *p* = 0.5 and added zero-mean Gaussian noise (*σ* = 0.4) before binarising. To quantify the significance value of the interactions, we generated 1,000 bootstrap resamples of the data, and calculated *F* : the fraction of resampled interactions that have a different sign from the original interaction. A smaller *F* corresponds to a more significant interaction.

**Figure S3:**
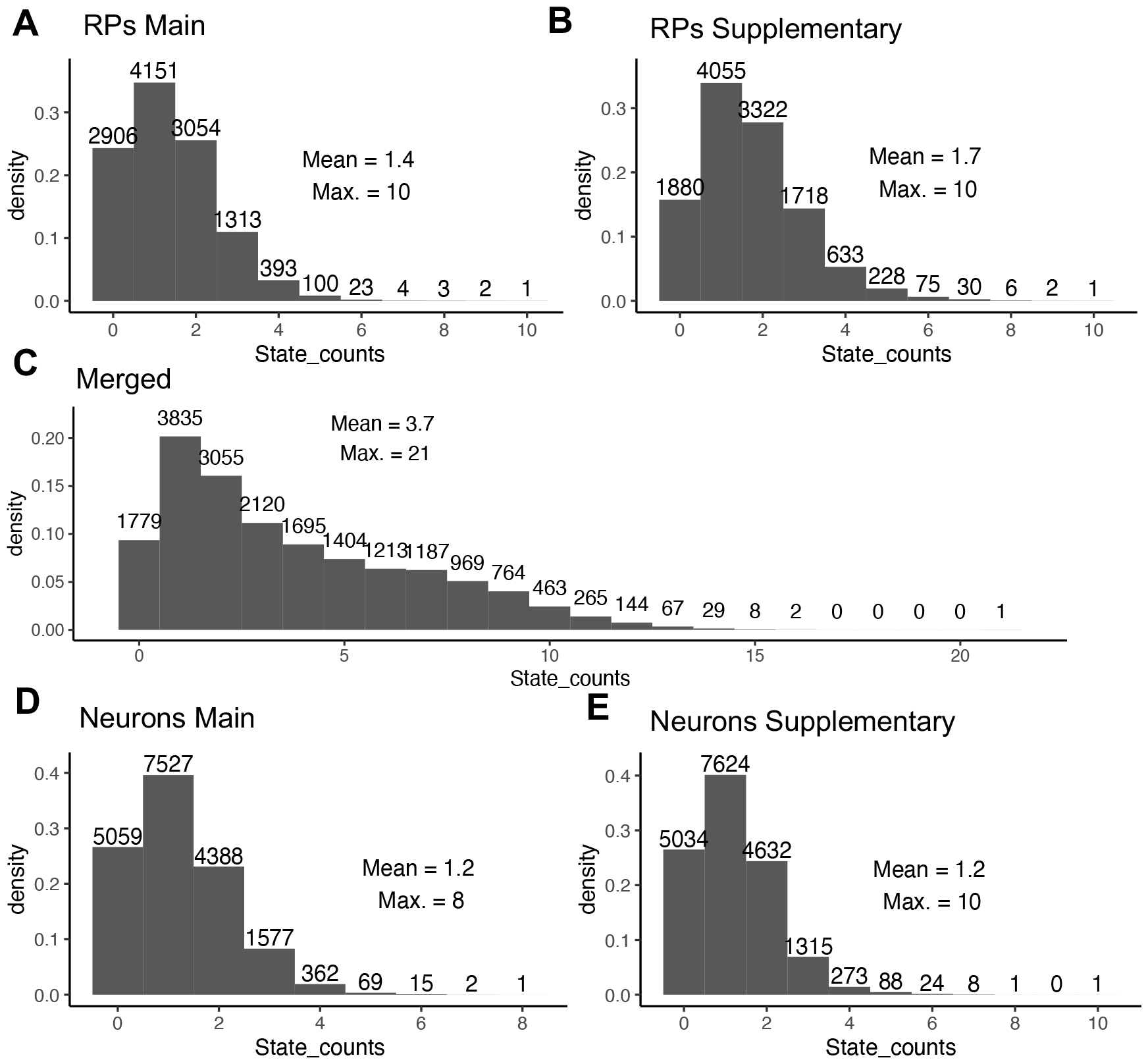
Histograms of the number of cells labelled with variable numbers of states. Results are for two disjoint embryonic radial glial cell-like cells (A and B), for the merged RP and neuron set (C), and for two disjoint sets of developmental neurons (D and E). Numbers shown above the bars indicate how the number of cells with the specific number of labels (X-axis).

### 7.1. Nextflow runtime on HPC cluster

Stator’s nextflow pipeline is represented in Fig. S4.

**Figure S4:**
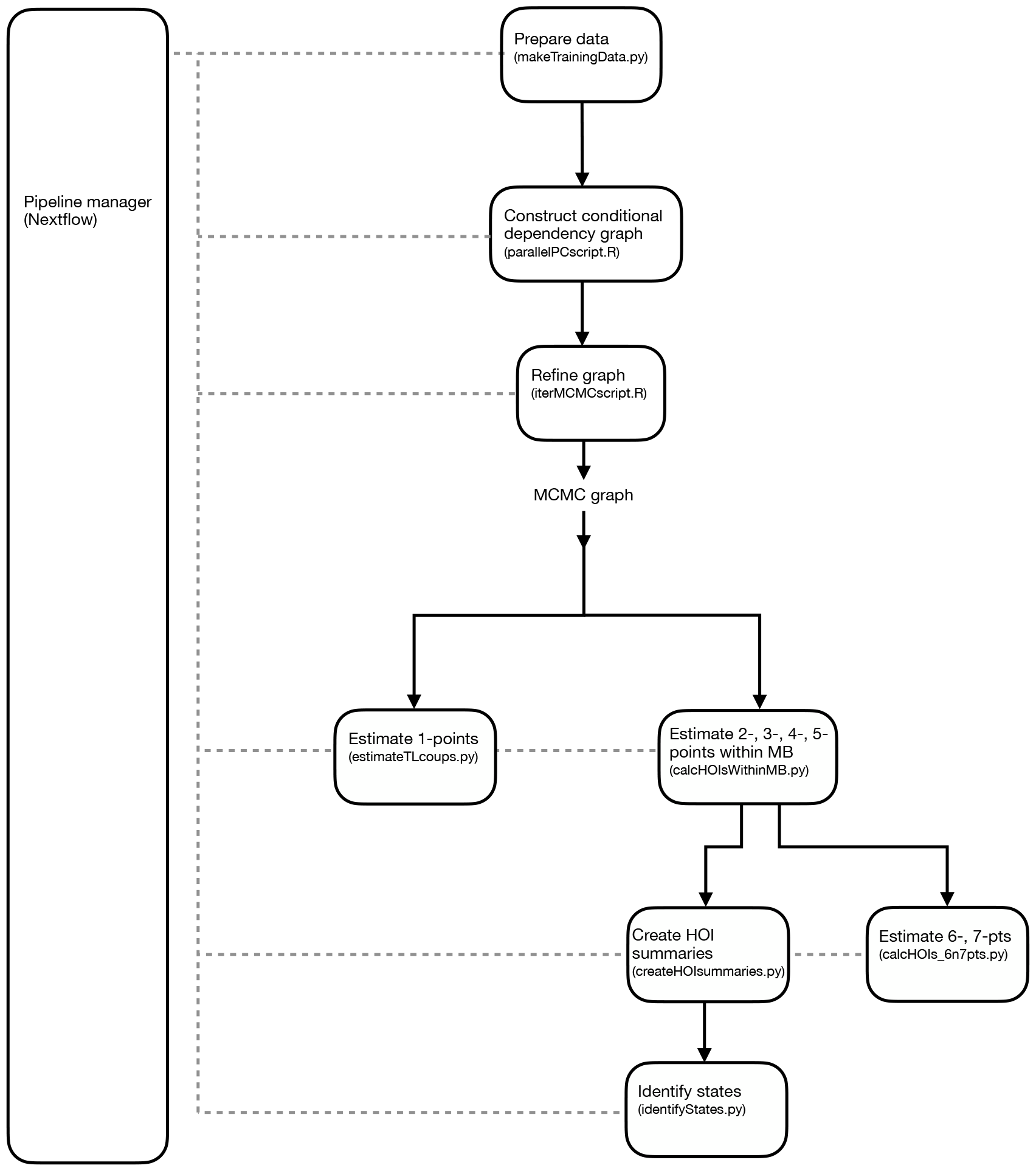
Stator’s nextflow pipeline. From data preparation to groupings of d-tuples used for state identification. Abbreviations: HOI, Higher-order interactions; MB, Markov blanket.

Stator computes combinatorial 2-point up to 7-point gene interactions, if estimable. The complete set of interactions among 1,000 HVGs is not computationally tractable: half a million 2-point interaction (1,000 choose 2), 166 million 3-point interactions, 41 billion 4-point interactions, 8 trillion 5-point interactions and so on. Instead, Stator first uses the Peter-Clark constraint-based algorithm (Spirtes et al., 2001), in order to perform (conditional) independence tests in order of increasing complexity, starting from a fully connected (undirected) graph of genes. Once a primary skeleton is obtained via the PC algorithm, Stator then uses a score based MCMC algorithm to improve the accuracy of an estimated Directed Acyclic Graph (DAG), using the hybrid approach introduced in Kuipers et al. (2022). This massively reduces the search space for potentially significant interactions. Note that memory requirements and runtime depend on gene expression connectivity in the given tissue and condition of interest. The sparser the dependence structure, the lower are the memory and runtime requirements which are unknown prior to running Stator’s nextflow pipeline. For a mouse embryonic brain dataset with 19,000 cells, the maximum memory requirement was 32G for running the nextflow pipeline.

**Figure S5:**
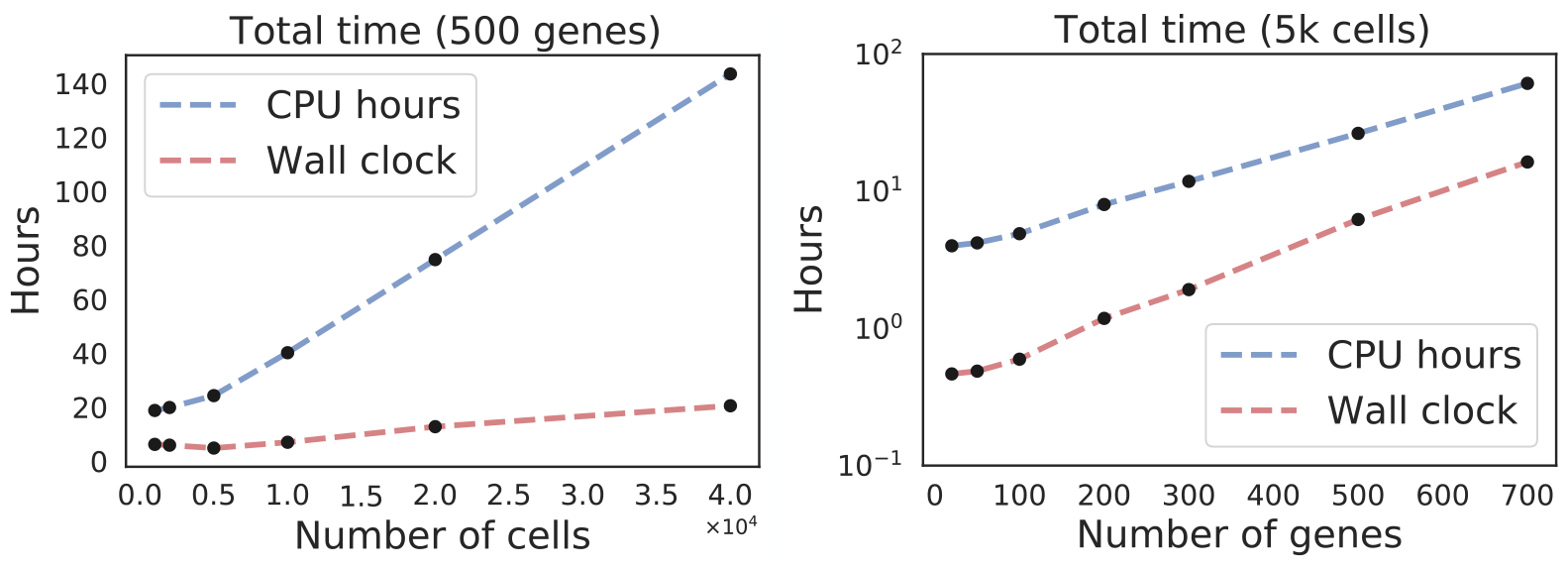
Runtime of Stator’s nextflow pipeline for increasing numbers of cells (left) or genes (right), using a fixed set of HPC resource. Runtime is linear in the number of cells and exponential in the number of genes. The ‘wall clock’ time in red shows the total amount of time lapsed between the beginning and end of the nextflow process, including the scheduling time on the HPC, and corresponds to the real time spent to obtain results. CPU time includes only the hours taken for tasks spent on the CPU. However, due to parallelisation, the CPU time will be longer than the wall clock. The wall clock depends on availability of HPC parallelisation resources. For example, for the 10X data set with 19,000 cells and 1,000 genes, the run time was approximately 5 days on the Edinburgh University High Performance Compute Cluster (Eddie), parallelised over multiple cores corresponding to 2,500 CPU hours, as reported by the Nextflow pipeline manager.

**Figure S6:**
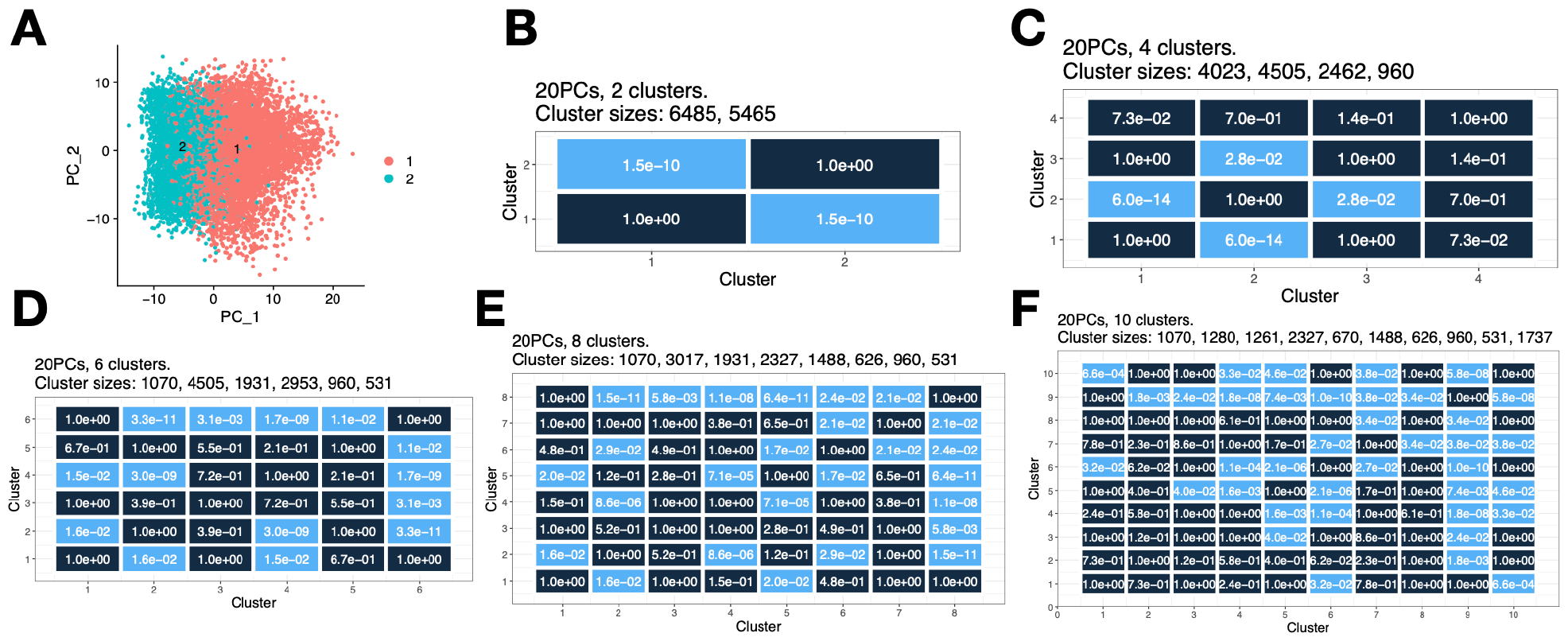
Embryonic radial glial cell precursor cell hierarchical clustering analysis, with p-value quantification bounding for selective inference (Gao et al., 2022). We applied standard single cell clustering to the non-binarised data, followed by pair-wise p-value quantification of cluster significance (Gao et al., 2022). In panels B-F, light blue represents pairwise significantly distinct clusters with Bonferroni corrected p-value *<* 0.05, whereas dark blue represents not significantly distinct clusters. **(A)** According to the clustering analysis this dataset is relatively homogeneous, containing two significantly distinct clusters, shown in a PCA plot. **(B)** Heatmap showing the p-value between clusters when choosing *k* = 2 clusters. **(C-D)** Heatmaps showing statistical support for clusters when choosing *k* = 4, 6, 8, 10 (C, D, E, F). Under these scenarios, not every cluster pair is significantly separated (*p <* 0.05 after correction with the Bonferroni method). To declare the final number of distinct clusters, we take the largest number of clusters such that all clusters are pairwise significantly distinct as *k* changes. In this case, it can be observed from panels B and C, that clusters #1, 2 are significantly pairwise distinct, but clusters #3, 4 are not. Therefore, we declare *k* = 2 distinct clusters.

**Figure S7:**
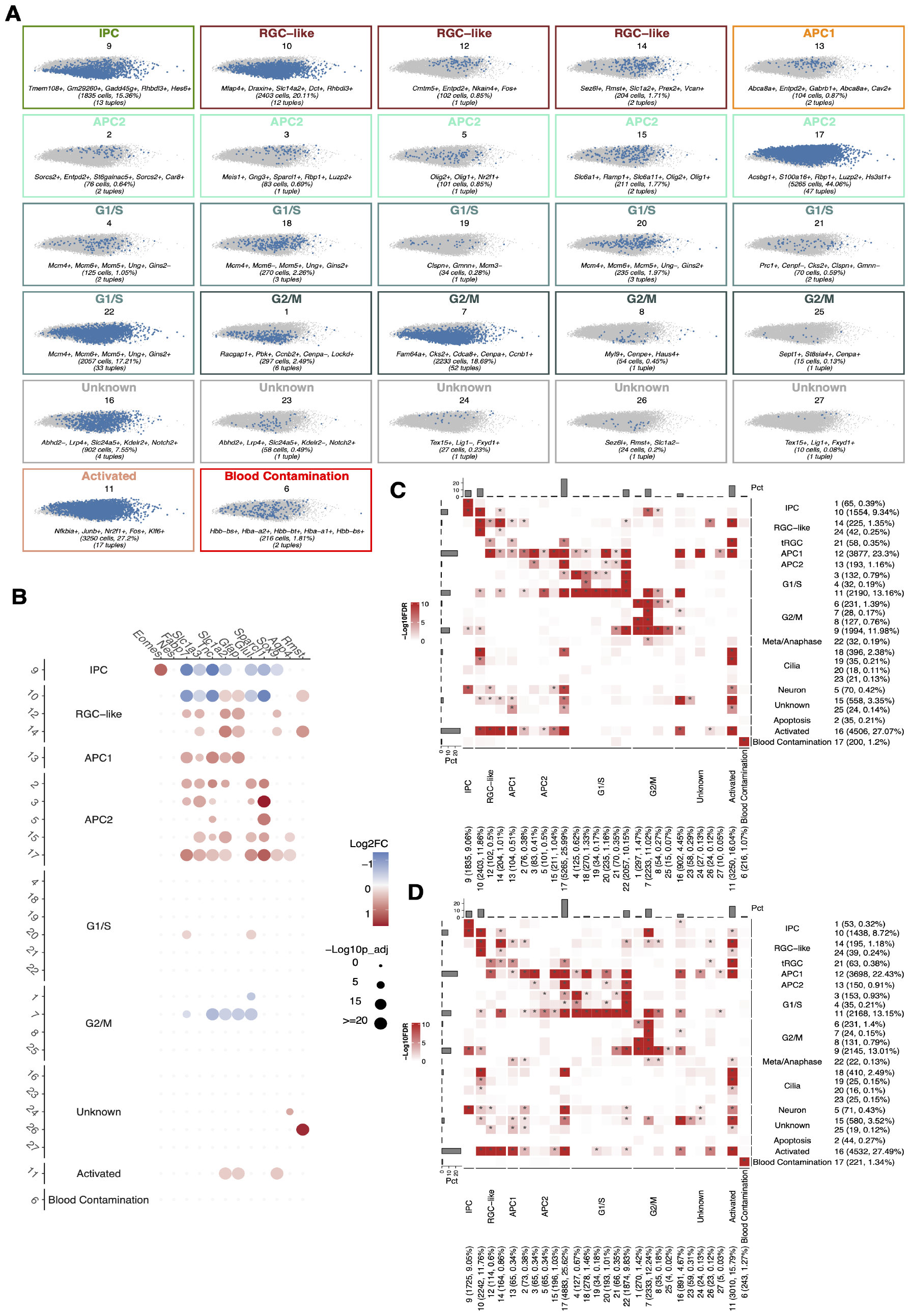
Extended Figure for Fig. 2. Disjoint RP dataset tested for reproducibility. Figure S7: **(A)** Stator identifies 27 states for a disjoint *N* = 11, 950 embryonic radial glial-like cells at maximum modularity, annotated on the basis of d-tuple and s2o-DE genes as markers for cell types or cell cycle phases. **(B)** Dot plot illustrating differential expression of astrocytogenesis marker genes across all 27 Stator states. The size of the dots represents the -log_10_(Seurat p-val-adj) from differential gene expression testing between a state and all other states. Colour intensity reflects the log_2_(FC) of gene expression. **(C)** Heatmap showing the significance of d-tuple enrichment for d-tuples shown in Fig. 2 (Y-axis) among cells represented in panel A (X-axis). Significance was assessed using the hypergeometric test. **(D)** As C, but the significance of d-tuple enrichment for d-tuples shown in panel A among cells represented in Fig. 2. These two disjoint datasets show a good degree of replication.

**Figure S8:**
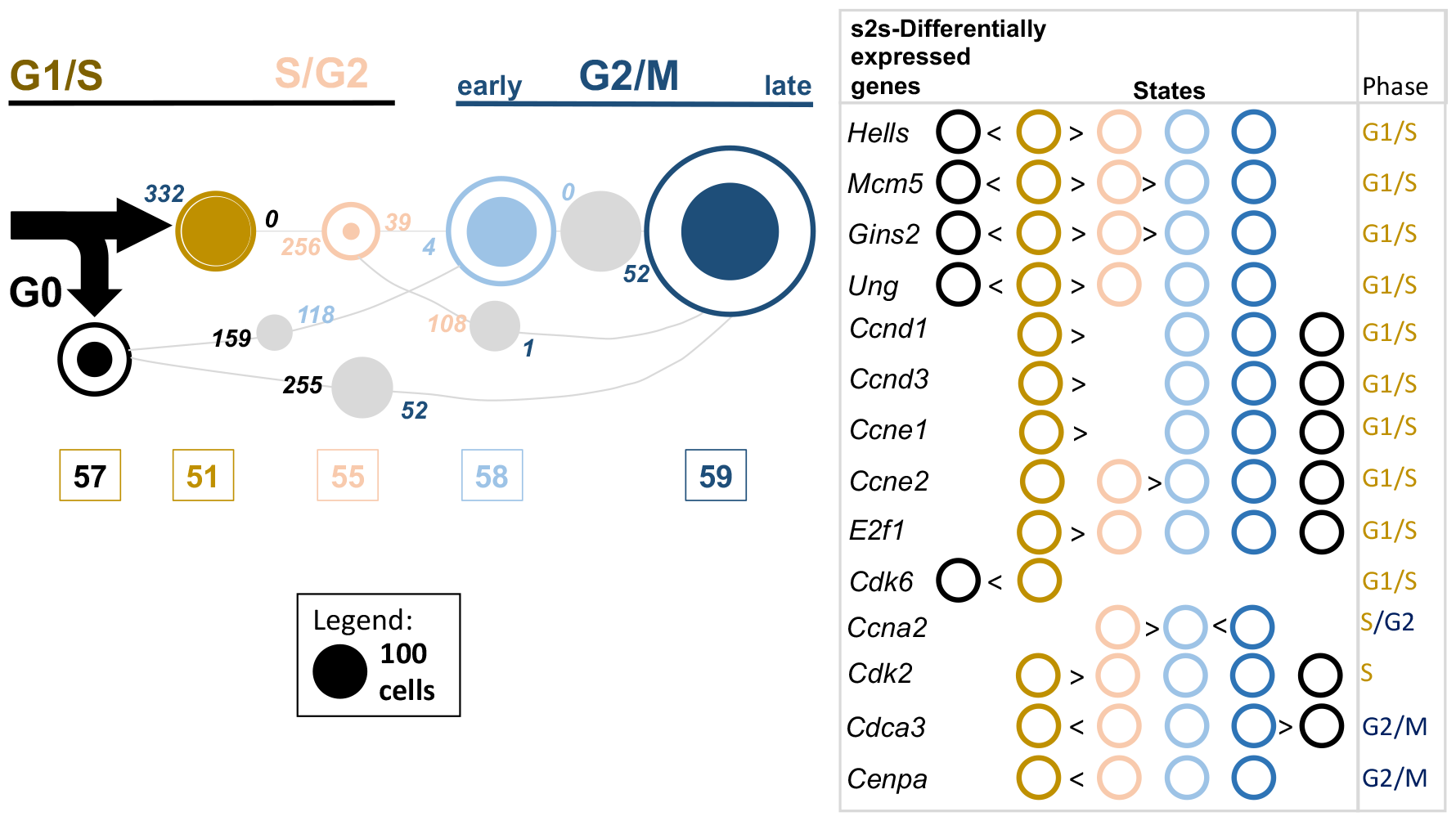
Extended Figure 3. s2s-DEG analysis for 5 cell cycle states. Numbers of cells labelled with any one of 5 cell cycle states (#57, 51, 55, 58, and 59) in embryonic RPs and neurons; areas of circles are proportional to their number (see legend). Filled circles indicate numbers of cells labelled with only of these single cell cycle states. Grey circles’ areas indicate numbers of cells labelled with two cell cycle states, those indicated by lines. Numbers of significantly differentially expressed genes between cell cycle state pairs (*i.e*., s2s-DEGs) are provided between the two states being compared; their colours refer to the state showing higher expression. State pairs with ≤ 25 cells are not shown. Right: s2s-DEGs are indicated by “*>*“ or “*<*“ symbols; for example, *Hells* mRNA expression is significantly higher in State #51 over States #57, 55, 58 and 59. Early/late G1/S or G2/M cell cycle phase labels (top) were assigned using these mRNAs’ cell cycle phases known from high-throughput (top right; (Giotti et al., 2018)) and targeted experiments (*Ung* mRNA in late G1/S (Slupphaug et al., 1991) and *Cenpa* in G2 (Shelby et al., 1997)).

**Figure S9:**
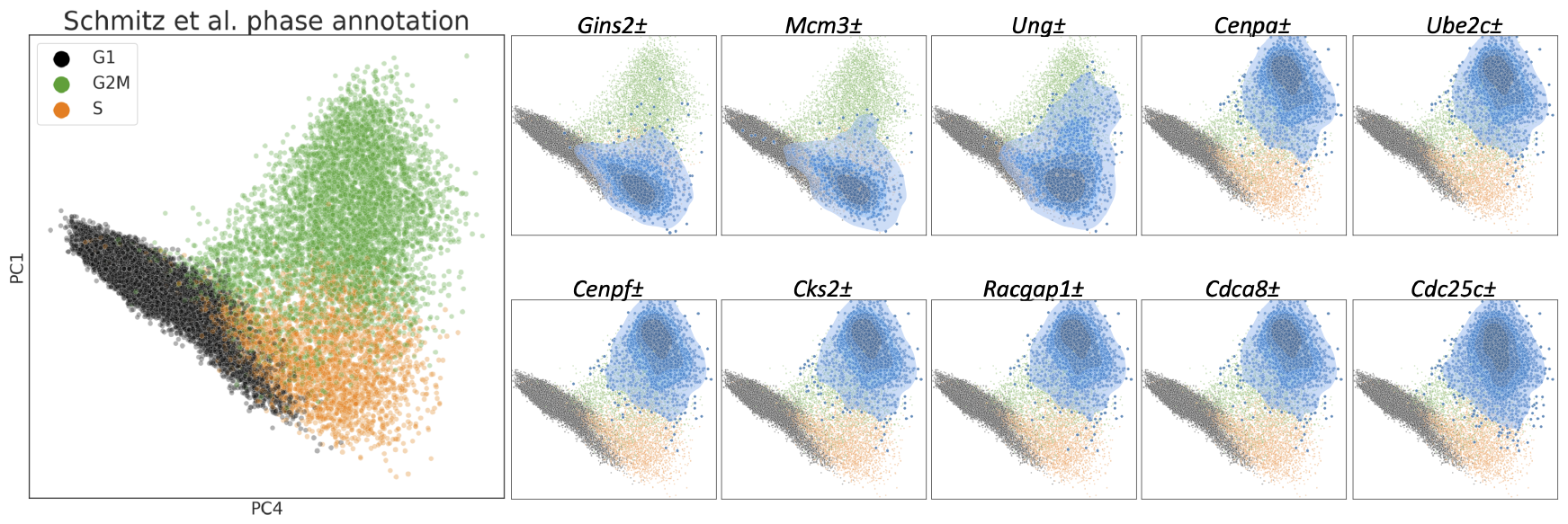
Extended Figure 3. *Minus* gene expression is required to specify cell cycle subphases. Left: External cell cycle annotations of a merged mouse brain data set sourced from five different experiments (Schmitz et al., 2022), with the cells from data set GSE93421 removed. Right: In contrast to Fig. 3C, highlighting cells based only on the expression of all but one of the marker genes, leaving the gene indicated above each plot (the ‘minus’ gene) unrestricted, results in more diffuse cell cycle specificity, and simply marks cells in G1/S or G2M phases.

**Figure S10:**
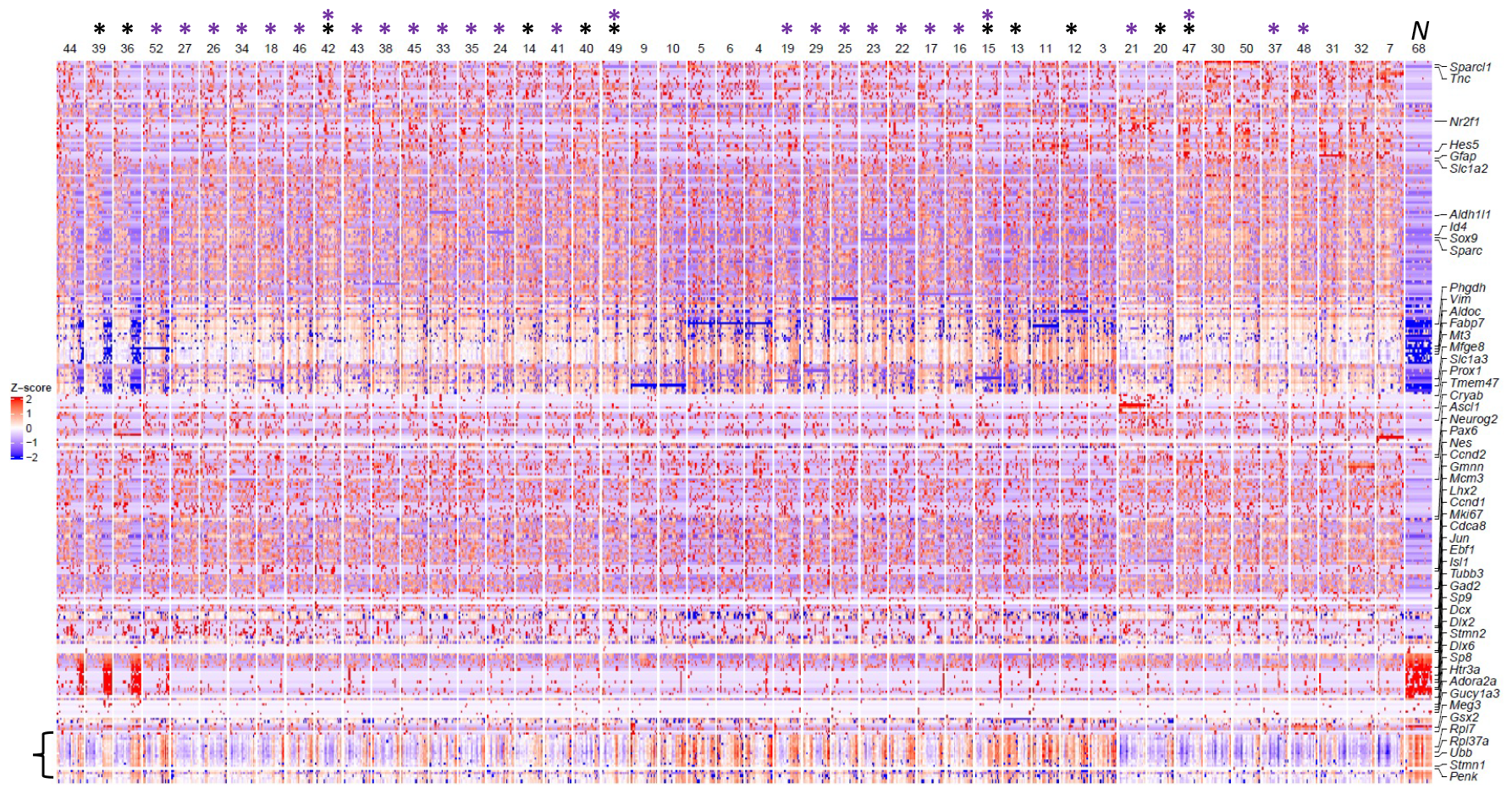
Heatmap of unbinarised single cell gene expression across 47 embryonic radial glial precursor cell (eRP) states and, for comparison, a single *Tubb3+* neuron state (State 68 [“N”]: *Slain1* [1], *Phgdh* [0], *Tubb3* [1]). Top: Asterisks indicate states with higher expression of neuronal marker s2s-DEGs than State 44 (black) or State 40 (purple). Numbers indicate States. Right: The 20 highest differentially expressed genes per state in s2oDEGs analysis, with, in addition, E17.5 eRP expressing genes listed in Supplementary Table S3 of Yuzwa et al. (2017) are shown. Horizontal lines reflect genes that are not expressed (blue) or always expressed (red) in a State’s cells. Heterogeneous gene expression across states is evident, for example, due to high expression of ribosomal protein and other genes (indicated by a bracket, lower left), an indicator of activated neural stem cells (Borrett et al., 2022; Dulken et al., 2017).

**Figure S11:**
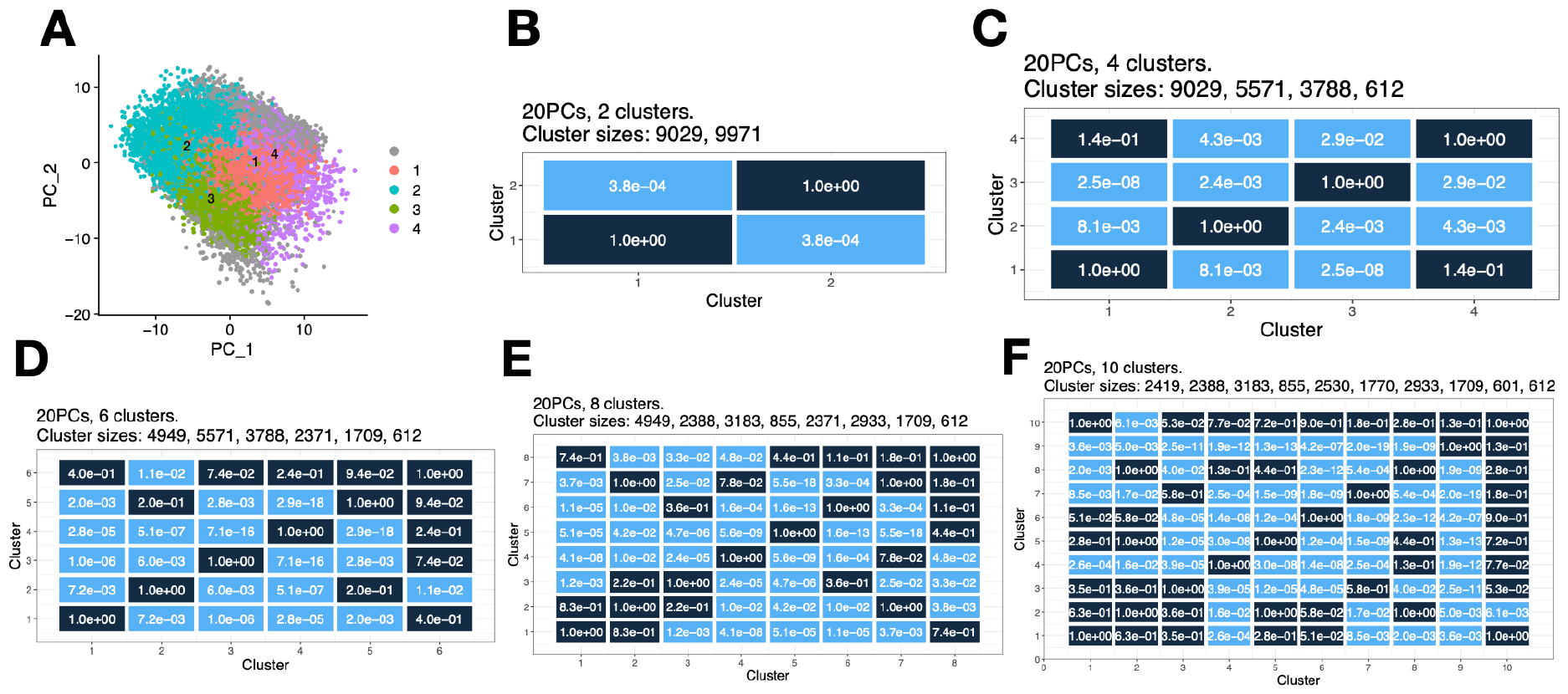
Hierarchical clustering of E18 developmental neurons, with p-value quantification bounding for selective inference (Gao et al., 2022). We applied standard single cell clustering to the non-binarised data, followed by pair-wise p-value quantification of cluster significance (Gao et al., 2022). In panels B-F, light blue represents pairwise significantly distinct clusters with Bonferroni correction p-value *<* 0.05, whereas dark blue represents not significantly distinct clusters. **(A)** PCA plot showing clustering of cells. According to this clustering analysis the dataset contains two significantly distinct clusters. **(B)** Heatmap showing the p-value for *k* = 2 clusters. It leads to significant differences between the clusters. **(C-F)** Heatmaps showing p-values between cluster pairs for *k* = 4, 6, 8, 10. Under these scenarios, not every cluster pair is significantly separated (*p <* 0.05 after correction with the Bonferroni method). To declare the final number of distinct clusters, we take the largest number of clusters such that all clusters are pairwise significantly distinct as *k* changes. In this case, it can be observed from panel D that clusters #1, 2, 3, 4 are significantly pairwise distinct, but not clusters #5, 6. Therefore, we declare *k* = 4 distinct clusters.

**Figure S12:**
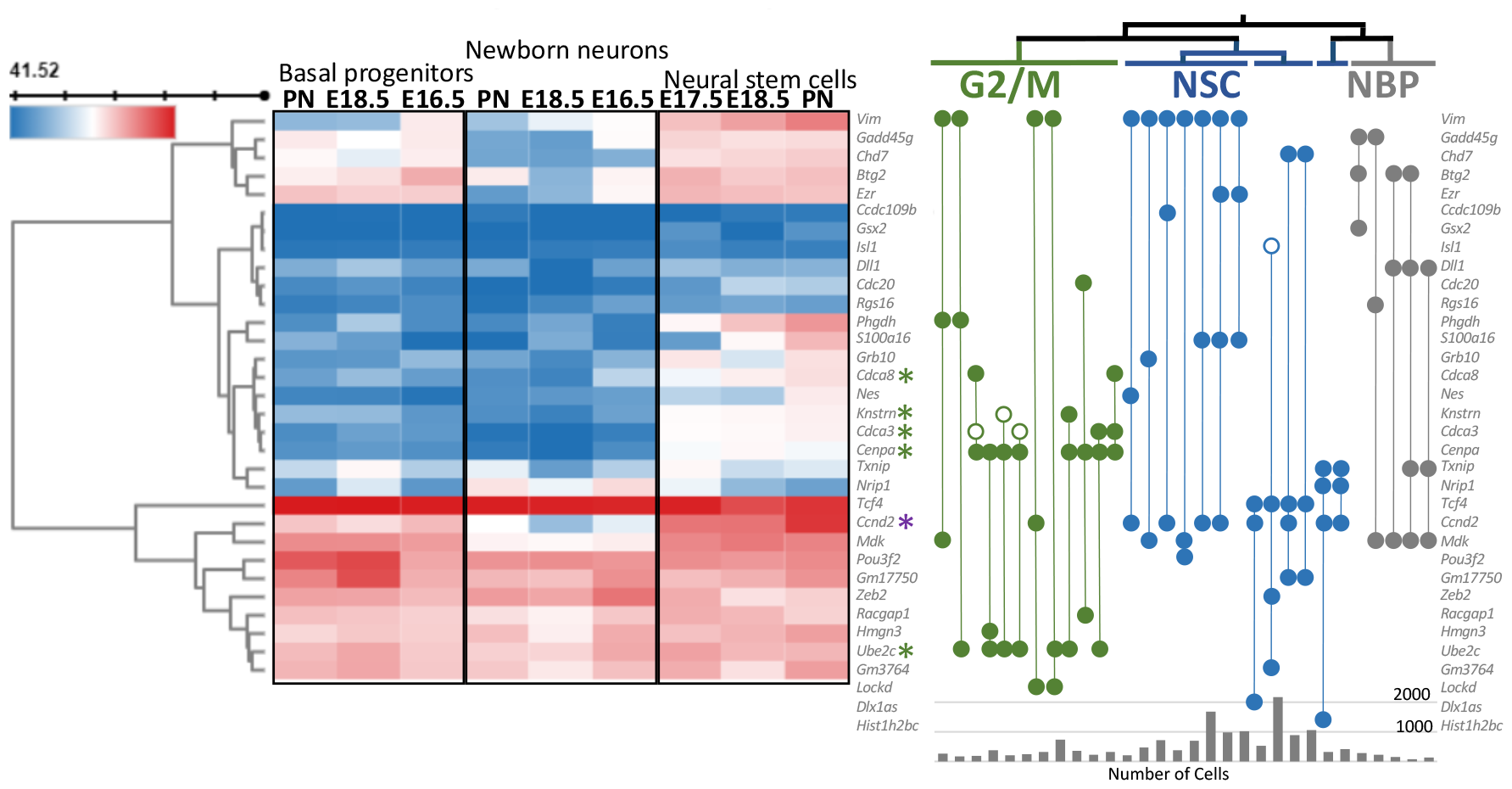
Example of a Stator state (#26), identified using a high Dice similarity (0.94), which can be further separated into biological states, here G2/M cell cycle phase, neural stem cells (NSC) and newborn progenitor (NPC) cells. Differential expression across embryonic (E) day 16.5 to postnatal day 1 of genes specifying d-tuples in neuron developmental data set #26. The heatmap indicates gene expression in NSCs, basal progenitor cells and newborn neurons (NBN) taken from NeuroStemX, an external data set https://neurostemx.ethz.ch (Mukhtar et al., 2022); expression data for *Lockd, Dlx1as* and *Hist1h2bc* was not available. Based on literature markers for G2/M cell cycle phase (green asterisk), NSCs and NB progenitors (NBP; *Btg2* (Micheli et al., 2015); *Mdk* (Winkler and Yao, 2014)), #26 d-tuples were partitioned into G2/M, NSC and NB progenitor states (in green, blue and grey, respectively); tuples are indicated by vertical lines linking either three or four genes (filled circle signifies expressed, unfilled circle not expressed genes). High cyclin D2 (*Ccnd2* ; purple asterisk) mRNA levels separates self-renewing cells from NBNs (Tsunekawa et al., 2014); *Ccnd2* -/- mice lack newly born neurons (Kowalczyk et al., 2004). Three tuples containing *Dll1* are notable in specifying newborn progenitor cells. *Dll1* is known to segregate asymmetrically during mitosis of NSCs, subsequently being inherited by differentiating cells (Kawaguchi et al., 2013).

**Figure S13:**
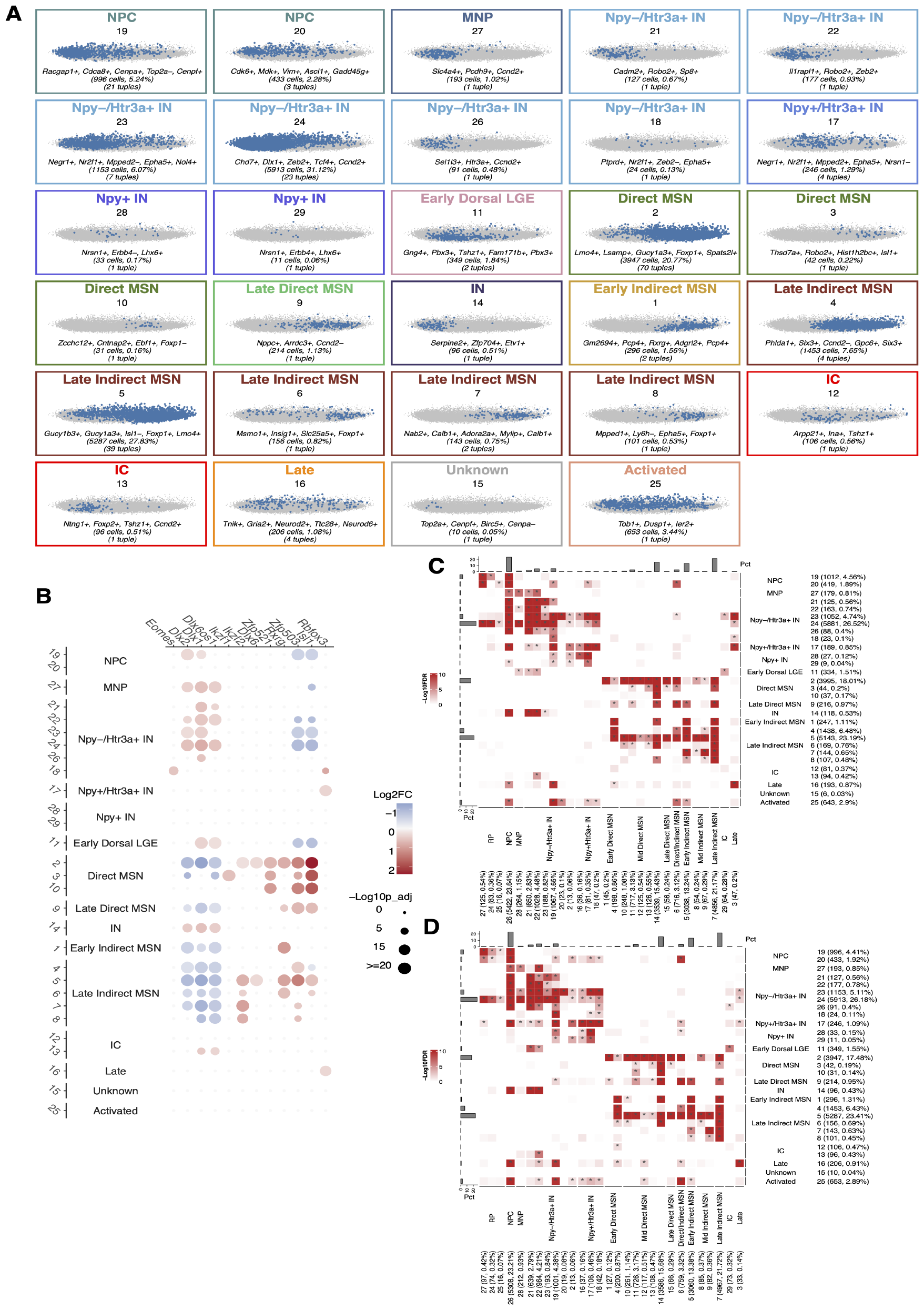
Extended Figure 4. A second dataset of developmental neurons testing for reproducibility. Disjoint neurons dataset for reproducibility. **(A)** Stator identifies 29 signatures at a Dice similarity of 0.94, annotated when d-tuple and/or s2o-DEGs are marker genes for cell (sub)types or states. **(B)** Dot plot illustrating differential expression of neurogenesis marker genes across all Stator states. The size of the dots represents the -log_10_(Seurat p-val-adj) from differential expression testing between a state and all other states. Colour intensity represents the log_2_(FC) of gene expression. **(C)** Heatmap showing the significance of d-tuple enrichment for d-tuples shown in Fig. 4 (Y-axis) among cells represented in panel A (X-axis). Significance was assessed using the hypergeometric test and Benjamini-Hochberg procedure was applied to calculate the False Discovery Rate (FDR). **(D)** As C, but the significance of d-tuple enrichment for d-tuples shown in panel A among cells represented in Fig. 4. These two disjoint datasets show a good degree of replication.

**Figure S14:**
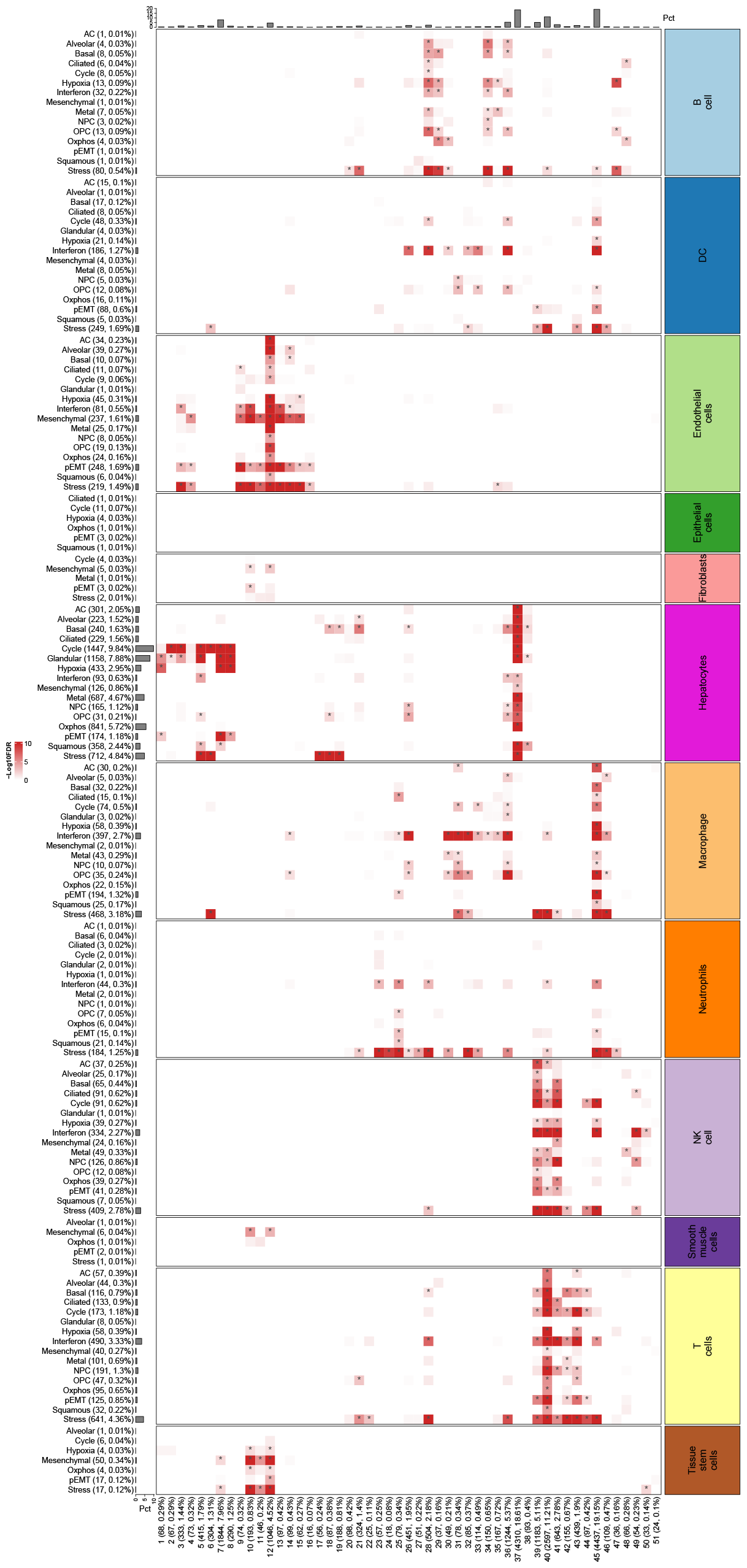
Extended Figure 6. Stator states (columns) that are significantly enriched in HCC cells previously doubly-annotated by cell type (right) and cell state (left) inferred using singleR and NMF, respectively.

## Notes

### Competing Interest Statement

The authors have declared no competing interest.

## References

10x Genomics (2017). Transcriptional profiling of 1.3 million brain cells with the chromium single cell 3’solution.

Akdemir, E., Huang, A., and Deneen, B. (2020). Astrocytogenesis: where, when, and how [version 1; peer review: 2 approved]. F1000Research, 9(233).

Aleksander, S. A., Balhoff, J., Carbon, S., Cherry, J. M., Drabkin, H. J., Ebert, D., Feuermann, M., Gaudet, P., Harris, N. L., et al. (2023). The Gene Ontology knowledgebase in 2023. Genetics, 224(1):iyad031.

Andrews, T. S., Atif, J., Liu, J. C., Perciani, C. T., Ma, X.-Z., Thoeni, C., Slyper, M., Eraslan, G., Segerstolpe, A., Manuel, J., et al. (2022). Single-cell, single-nucleus, and spatial rna sequencing of the human liver identifies cholangiocyte and mesenchymal heterogeneity. Hepatol Commun, 6(4):821–840.

Antebi, Y. E., Linton, J. M., Klumpe, H., Bintu, B., Gong, M., Su, C., McCardell, R., and Elowitz, M. B. (2017). Combinatorial signal perception in the bmp pathway. Cell, 170(6):1184–1196.

Aran, D., Looney, A. P., Liu, L., Wu, E., Fong, V., Hsu, A., Chak, S., Naikawadi, R. P., Wolters, P. J., Abate, A. R., et al. (2019). Reference-based analysis of lung single-cell sequencing reveals a transitional profibrotic macrophage. Nature Immunology, 20(2):163–172.

Arlotta, P., Molyneaux, B. J., Jabaudon, D., Yoshida, Y., and Macklis, J. D. (2008). Ctip2 controls the differentiation of medium spiny neurons and the establishment of the cellular architecture of the striatum. Journal of Neuroscience, 28(3):622–632.

Arnosti, D. N., Barolo, S., Levine, M., and Small, S. (1996). The eve stripe 2 enhancer employs multiple modes of transcriptional synergy. Development, 122(1):205–214.

Asahina, H., Masuba, A., Hirano, S., and Yuri, K. (2012). Distribution of protocadherin 9 protein in the developing mouse nervous system. Neuroscience, 225:88–104.

Ashburner, M., Ball, C. A., Blake, J. A., Botstein, D., Butler, H., Cherry, J. M., Davis, A. P., Dolinski, K., Dwight, S. S., Eppig, J. T., et al. (2000). Gene ontology: tool for the unification of biology. Nature Genetics, 25(1):25–29.

Barkley, D., Moncada, R., Pour, M., Liberman, D. A., Dryg, I., Werba, G., Wang, W., Baron, M., Rao, A., Xia, B., et al. (2022). Cancer cell states recur across tumor types and form specific interactions with the tumor microenvironment. Nature Genetics, 54(8):1192–1201.

Beentjes, S. V. and Khamseh, A. (2020). Higher-order interactions in statistical physics and machine learning: A model-independent solution to the inverse problem at equilibrium. Phys. Rev. E, 102:053314.

Benjamini, Y. and Hochberg, Y. (1995). Controlling the false discovery rate: A practical and powerful approach to multiple testing. Journal of the Royal Statistical Society. Series B (Methodological), 57(1):289–300.

Benjamini, Y. and Yekutieli, D. (2001). The control of the false discovery rate in multiple testing under dependency. The Annals of Statistics, 29(4):1165–1188.

Biagioli, M., Pinto, M., Cesselli, D., Zaninello, M., Lazarevic, D., Roncaglia, P., Simone, R., Vlachouli, C., Plessy, C., Bertin, N., et al. (2009). Unexpected expression of α- and β-globin in mesencephalic dopaminergic neurons and glial cells. Proceedings of the National Academy of Sciences, 106(36):15454–15459.

Blondel, V. D., Guillaume, J.-L., Lambiotte, R., and Lefebvre, E. (2008). Fast unfolding of communities in large networks. Journal of statistical mechanics: theory and experiment, 2008(10):P10008.

Borrett, M. J., Innes, B. T., Tahmasian, N., Bader, G. D., Kaplan, D. R., and Miller, F. D. (2022). A shared transcriptional identity for forebrain and dentate gyrus neural stem cells from embryogenesis to adulthood. eNeuro, 9(1).

Bouland, G. A., Mahfouz, A., and Reinders, M. J. T. (2021). Differential analysis of binarized single-cell RNA sequencing data captures biological variation. NAR Genomics and Bioinformatics, 3(4). lqab118.

Bouland, G. A., Mahfouz, A., and Reinders, M. J. T. (2023). Consequences and opportunities arising due to sparser single-cell rna-seq datasets. Genome Biology, 24(1):86.

Chari, T. and Pachter, L. (2023). The specious art of single-cell genomics. PLOS Computational Biology, 19(8):1–20.

Cirnaru, M.-D., Song, S., Tshilenge, K.-T., Corwin, C., Mleczko, J., Aguirre, C. G., Benlhabib, H., Bendl, J., Apontes, P., Fullard, J., et al. (2021). Unbiased identification of novel transcription factors in striatal compartmentation and striosome maturation. eLife, 10:e65979.

Colombo, D., Maathuis, M. H., et al. (2014). Order-independent constraint-based causal structure learning. J. Mach. Learn. Res., 15(1):3741–3782.

Coré, N., Erni, A., Hoffmann, H. M., Mellon, P. L., Saurin, A. J., Beclin, C., and Cremer, H. (2020). Stem cell regionalization during olfactory bulb neurogenesis depends on regulatory interactions between Vax1 and Pax6. eLife, 9:e58215.

Csardi, G., Nepusz, T., et al. (2006). The igraph software package for complex network research. InterJournal, complex systems, 1695(5):1–9.

Cui, G., Jun, S. B., Jin, X., Pham, M. D., Vogel, S. S., Lovinger, D. M., and Costa, R. M. (2013). Concurrent activation of striatal direct and indirect pathways during action initiation. Nature, 494(7436):238–242.

Dann, E., Cujba, A.-M., Oliver, A. J., Meyer, K. B., Teichmann, S. A., and Marioni, J. C. (2023). Precise identification of cell states altered in disease using healthy single-cell references. Nature Genetics.

Dann, E., Henderson, N. C., Teichmann, S. A., Morgan, M. D., and Marioni, J. C. (2022). Differential abundance testing on single-cell data using k-nearest neighbor graphs. Nature Biotechnology, 40(2):245–253.

Dulken, B. W., Leeman, D. S., Boutet, S. C., Hebestreit, K., and Brunet, A. (2017). Single-cell transcriptomic analysis defines heterogeneity and transcriptional dynamics in the adult neural stem cell lineage. Cell Reports, 18(3):777–790.

Efron, B. (1979). Computers and the theory of statistics: Thinking the unthinkable. SIAM Review, 21(4):460–480.

Ericson, J., Rashbass, P., Schedl, A., Brenner-Morton, S., Kawakami, A., van Heyningen, V., Jessell, T. M., and Briscoe, J. (1997). Pax6 controls progenitor cell identity and neuronal fate in response to graded shh signaling. Cell, 90(1):169–180.

Fischer, M., Grossmann, P., Padi, M., and DeCaprio, J. A. (2016). Integration of TP53, DREAM, MMB-FOXM1 and RB-E2F target gene analyses identifies cell cycle gene regulatory networks. Nucleic Acids Research, 44(13):6070–6086.

Franzén, O., Gan, L.-M., and Björkegren, J. L. M. (2019). PanglaoDB: a web server for exploration of mouse and human single-cell RNA sequencing data. Database, 2019:baz046.

Fuccillo, M. V., Földy, C., Gökce, Ö., Rothwell, P. E., Sun, G. L., Malenka, R. C., and Südhof, T. C. (2015). Single-cell mrna profiling reveals cell-type-specific expression of neurexin isoforms. Neuron, 87(2):326–340.

Gao, L. L., Bien, J., and Witten, D. (2022). Selective inference for hierarchical clustering. Journal of the American Statistical Association, pages 1–11.

Gaujoux, R. and Seoighe, C. (2010). A flexible r package for nonnegative matrix factorization. BMC Bioinformatics, 11(1):367.

Ghazanfar, S., Lin, Y., Su, X., Lin, D. M., Patrick, E., Han, Z.-G., Marioni, J. C., and Yang, J. Y. H. (2020). Investigating higher-order interactions in single-cell data with schot. Nature methods, 17(8):799–806.

Giotti, B., Chen, S.-H., Barnett, M. W., Regan, T., Ly, T., Wiemann, S., Hume, D. A., and Freeman, T. C. (2018). Assembly of a parts list of the human mitotic cell cycle machinery. Journal of Molecular Cell Biology, 11(8):703–718.

Götz, M., Sirko, S., Beckers, J., and Irmler, M. (2015). Reactive astrocytes as neural stem or progenitor cells: In vivo lineage, in vitro potential, and genome-wide expression analysis. Glia, 63(8):1452–1468.

Gu, Z. (2022). Complex heatmap visualization. iMeta, 1(3):e43.

Gu, Z., Eils, R., and Schlesner, M. (2016). Complex heatmaps reveal patterns and correlations in multidimensional genomic data. Bioinformatics, 32(18):2847–2849.

Hernan, M. and Robins, J. (2023). Causal Inference. Chapman & Hall/CRC Monographs on Statistics & Applied Probab. CRC Press.

Imayoshi, I. and Kageyama, R. (2014). bhlh factors in self-renewal, multipotency, and fate choice of neural progenitor cells. Neuron, 82(1):9–23.

Jansma, A. (2023a). Higher-order interactions and their duals reveal synergy and logical dependence beyond shannon-information. Entropy, 25(4):648.

Jansma, A. (2023b). Higher-order interactions in single-cell gene expression. PhD thesis, University of Edinburgh.

Kawaguchi, D., Furutachi, S., Kawai, H., Hozumi, K., and Gotoh, Y. (2013). Dll1 maintains quiescence of adult neural stem cells and segregates asymmetrically during mitosis. Nature Communications, 4(1):1880.

Kawauchi, T., Chihama, K., Nabeshima, Y.-i., and Hoshino, M. (2003). The in vivo roles of stef/tiam1, rac1 and jnk in cortical neuronal migration. The EMBO Journal, 22(16):4190–4201.

Kotliar, D., Veres, A., Nagy, M. A., Tabrizi, S., Hodis, E., Melton, D. A., and Sabeti, P. C. (2019). Identifying gene expression programs of cell-type identity and cellular activity with single-cell rna-seq. Elife, 8:e43803.

Kowalczyk, A., Filipkowski, R. K., Rylski, M., Wilczynski, G. M., Konopacki, F. A., Jaworski, J., Ciemerych, M. A., Sicinski, P., and Kaczmarek, L. (2004). The critical role of cyclin D2 in adult neurogenesis. Journal of Cell Biology, 167(2):209–213.

Kuerbitz, J., Arnett, M., Ehrman, S., Williams, M. T., Vorhees, C. V., Fisher, S. E., Garratt, A. N., Muglia, L. J., Waclaw, R. R., and Campbell, K. (2018). Loss of intercalated cells (itcs) in the mouse amygdala of tshz1 mutants correlates with fear, depression, and social interaction phenotypes. Journal of Neuroscience, 38(5):1160–1177.

Kuipers, J., Suter, P., and Moffa, G. (2022). Efficient sampling and structure learning of Bayesian networks. Journal of Computational and Graphical Statistics, 31(3):639–650.

Kuzmin, E., VanderSluis, B., Wang, W., Tan, G., Deshpande, R., Chen, Y., Usaj, M., Balint, A., Mattiazzi Usaj, M., Van Leeuwen, J., et al. (2018). Systematic analysis of complex genetic interactions. Science, 360(6386):eaao1729.

Lacar, B., Linker, S. B., Jaeger, B. N., Krishnaswami, S. R., Barron, J. J., Kelder, M. J. E., Parylak, S. L., Paquola, A. M., Venepally, P., Novotny, M., O’Connor, C., Fitzpatrick, C., Erwin, J. A., Hsu, J. Y., Husband, D., McConnell, M. J., Lasken, R., and Gage, F. H. (2016). Nuclear rna-seq of single neurons reveals molecular signatures of activation. Nature Communications, 7(1):11022.

Le, T. D., Hoang, T., Li, J., Liu, L., Liu, H., and Hu, S. (2016). A fast pc algorithm for high dimensional causal discovery with multi-core pcs. IEEE/ACM transactions on computational biology and bioinformatics, 16(5):1483–1495.

Leiper, K., Croll, A., Booth, N. A., Moore, N. R., Sinclair, T., and Bennett, B. (1994). Tissue plasminogen activator, plasminogen activator inhibitors, and activator-inhibitor complex in liver disease. Journal of Clinical Pathology, 47(3):214–217.

Li, R. and Quon, G. (2019). scbfa: modeling detection patterns to mitigate technical noise in large-scale single-cell genomics data. Genome biology, 20:1–20.

Li, Z., Shang, Z., Sun, M., Jiang, X., Tian, Y., Yang, L., Wang, Z., Su, Z., Liu, G., li, X., You, Y., Yang, Z., Xu, Z., and Zhang, Z. (2022). Transcription factor sp9 is a negative regulator of d1-type msn development. Cell Death Discovery, 8(1):301.

Liu, J., Wu, X., and Lu, Q. (2022). Molecular divergence of mammalian astrocyte progenitor cells at early gliogenesis. Development, 149(5):dev199985.

Liu, J. K., Ghattas, I., Liu, S., Chen, S., and Rubenstein, J. L. (1997). Dlx genes encode dnabinding proteins that are expressed in an overlapping and sequential pattern during basal ganglia differentiation. Developmental Dynamics, 210(4):498–512.

Luecken, M. D. and Theis, F. J. (2019). Current best practices in single-cell rna-seq analysis: a tutorial. Molecular Systems Biology, 15(6):e8746.

Lun, A. T., McCarthy, D. J., and Marioni, J. C. (2016). A step-by-step workflow for low-level analysis of single-cell rna-seq data with bioconductor. F1000Research, 5.

MacParland, S. A., Liu, J. C., Ma, X.-Z., Innes, B. T., Bartczak, A. M., Gage, B. K., Manuel, J., Khuu, N., Echeverri, J., Linares, I., et al. (2018). Single cell rna sequencing of human liver reveals distinct intrahepatic macrophage populations. Nature Communications, 9(1):4383.

Micheli, L., Ceccarelli, M., Farioli-Vecchioli, S., and Tirone, F. (2015). Control of the normal and pathological development of neural stem and progenitor cells by the pc3/tis21/btg2 and btg1 genes. Journal of Cellular Physiology, 230(12):2881–2890.

Miyoshi, G., Young, A., Petros, T., Karayannis, T., Chang, M. M., Lavado, A., Iwano, T., Nakajima, M., Taniguchi, H., Huang, Z. J., Heintz, N., Oliver, G., Matsuzaki, F., Machold, R. P., and Fishell, G. (2015). Prox1 regulates the subtype-specific development of caudal ganglionic eminence-derived gabaergic cortical interneurons. Journal of Neuroscience, 35(37):12869–12889.

Morris, S. A. (2019). The evolving concept of cell identity in the single cell era. Development, 146(12):dev169748.

Mukhtar, T., Breda, J., Adam, M. A., Boareto, M., Grobecker, P., Karimaddini, Z., Grison, A., Eschbach, K., Chandrasekhar, R., Vermeul, S., et al. (2022). Temporal and sequential transcriptional dynamics define lineage shifts in corticogenesis. The EMBO Journal, 41(24):e111132.

Murtagh, F. and Legendre, P. (2014). Ward’s hierarchical agglomerative clustering method: which algorithms implement ward’s criterion? Journal of classification, 31:274–295.

Newman, M. E. (2006). Modularity and community structure in networks. Proceedings of the national academy of sciences, 103(23):8577–8582.

Parras, C. M., Schuurmans, C., Scardigli, R., Kim, J., Anderson, D. J., and Guillemot, F. (2002). Divergent functions of the proneural genes mash1 and ngn2 in the specification of neuronal subtype identity. Genes & Development, 16(3):324–338.

Pascual-Montano, A., Carazo, J., Kochi, K., Lehmann, D., and Pascual-Marqui, R. (2006). Non-smooth nonnegative matrix factorization (nsnmf). IEEE Transactions on Pattern Analysis and Machine Intelligence, 28(3):403–415.

Patel, A. P., Tirosh, I., Trombetta, J. J., Shalek, A. K., Gillespie, S. M., Wakimoto, H., Cahill, D. P., Nahed, B. V., Curry, W. T., Martuza, R. L., et al. (2014). Single-cell rna-seq highlights intratumoral heterogeneity in primary glioblastoma. Science, 344(6190):1396–1401.

Peters, C., He, S., Fermani, F., Lim, H., Ding, W., Mayer, C., and Klein, R. (2023). Transcriptomics reveals amygdala neuron regulation by fasting and ghrelin thereby promoting feeding. Science Advances, 9(21):eadf6521.

Petryniak, M. A., Potter, G. B., Rowitch, D. H., and Rubenstein, J. L. (2007). Dlx1 and dlx2 control neuronal versus oligodendroglial cell fate acquisition in the developing forebrain. Neuron, 55(3):417–433.

Poirier, K., Van Esch, H., Friocourt, G., Saillour, Y., Bahi, N., Backer, S., Souil, E., Castelnau-Ptakhine, L., Beldjord, C., Francis, F., Bienvenu, T., and Chelly, J. (2004). Neuroanatomical distribution of arx in brain and its localisation in gabaergic neurons. Molecular Brain Research, 122(1):35–46.

Ponting, C. P. (2019). The Human Cell Atlas: making ‘cell space’ for disease. Disease Models & Mechanisms, 12(2):dmm037622.

Precious, S., Kelly, C., Reddington, A., Vinh, N., Stickland, R., Pekarik, V., Scherf, C., Jeyasingham, R., Glasbey, J., Holeiter, M., Jones, L., Taylor, M., and Rosser, A. (2016). Foxp1 marks medium spiny neurons from precursors to maturity and is required for their differentiation. Experimental Neurology, 282:9–18.

Przysinda, A., Feng, W., and Li, G. (2020). Diversity of organism-wide and organ-specific endothelial cells. Current Cardiology Reports, 22(4):19.

Puram, S. V., Tirosh, I., Parikh, A. S., Patel, A. P., Yizhak, K., Gillespie, S., Rodman, C., Luo, C. L., Mroz, E. A., Emerick, K. S., et al. (2017). Single-cell transcriptomic analysis of primary and metastatic tumor ecosystems in head and neck cancer. Cell, 171(7):1611–1624.

Qiu, P. (2020). Embracing the dropouts in single-cell rna-seq analysis. Nature Communications, 11(1):1169.

Ramachandran, P., Dobie, R., Wilson-Kanamori, J., Dora, E., Henderson, B., Luu, N., Portman, J., Matchett, K., Brice, M., Marwick, J., et al. (2019). Resolving the fibrotic niche of human liver cirrhosis at single-cell level. Nature, 575(7783):512–518.

Regev, A., Teichmann, S. A., Lander, E. S., Amit, I., Benoist, C., Birney, E., Bodenmiller, B., Campbell, P., Carninci, P., Clatworthy, M., et al. (2017). Science forum: The human cell atlas. eLife, 6:e27041.

Revel, M., Sautes-Fridman, C., Fridman, W.-H., and Roumenina, L. T. (2022). C1q+ macrophages: passengers or drivers of cancer progression. Trends in Cancer, 8(7):517–526.

Riba, A., Oravecz, A., Durik, M., Jiménez, S., Alunni, V., Cerciat, M., Jung, M., Keime, C., Keyes, W. M., and Molina, N. (2022). Cell cycle gene regulation dynamics revealed by rna velocity and deep-learning. Nature Communications, 13(1):2865.

Ruan, X., Kang, B., Qi, C., Lin, W., Wang, J., and Zhang, X. (2021). Progenitor cell diversity in the developing mouse neocortex. Proceedings of the National Academy of Sciences, 118(10):e2018866118.

Rubenstein, J. and Campbell, K. (2020). Chapter 18 - neurogenesis in the basal ganglia. In Rubenstein, J., Rakic, P., Chen, B., and Kwan, K. Y., editors, Patterning and Cell Type Specification in the Developing CNS and PNS (Second Edition), pages 399–426. Academic Press, second edition edition.

Sapir, T., Levy, T., Sakakibara, A., Rabinkov, A., Miyata, T., and Reiner, O. (2013). Shootin1 acts in concert with kif20b to promote polarization of migrating neurons. Journal of Neuroscience, 33(29):11932–11948.

Saunders, A., Macosko, E. Z., Wysoker, A., Goldman, M., Krienen, F. M., de Rivera, H., Bien, E., Baum, M., Bortolin, L., Wang, S., et al. (2018). Molecular diversity and specializations among the cells of the adult mouse brain. Cell, 174(4):1015–1030.

Sayols, S. (2023). rrvgo: a bioconductor package for interpreting lists of gene ontology terms.

Schmitz, M. T., Sandoval, K., Chen, C. P., Mostajo-Radji, M. A., Seeley, W. W., Nowakowski, T. J., Ye, C. J., Paredes, M. F., and Pollen, A. A. (2022). The development and evolution of inhibitory neurons in primate cerebrum. Nature, 603(7903):871–877.

Shalek, A. K., Satija, R., Shuga, J., Trombetta, J. J., Gennert, D., Lu, D., Chen, P., Gertner, R. S., Gaublomme, J. T., Yosef, N., et al. (2014). Single-cell rna-seq reveals dynamic paracrine control of cellular variation. Nature, 510(7505):363–369.

Shang, Z., Yang, L., Wang, Z., Tian, Y., Gao, Y., Su, Z., Guo, R., Li, W., Liu, G., Li, X., Yang, Z., Li, Z., and Zhang, Z. (2022). The transcription factor zfp503 promotes the d1 msn identity and represses the d2 msn identity. Frontiers in Cell and Developmental Biology, 10.

Sharma, A., Seow, J. J. W., Dutertre, C.-A., Pai, R., Blériot, C., Mishra, A., Wong, R. M. M., Singh, G. S. N., Sudhagar, S., Khalilnezhad, S., et al. (2020). Onco-fetal reprogramming of endothelial cells drives immunosuppressive macrophages in hepatocellular carcinoma. Cell, 183(2):377–394.e21.

Shelby, R. D., Vafa, O., and Sullivan, K. F. (1997). Assembly of CENP-A into Centromeric Chromatin Requires a Cooperative Array of Nucleosomal DNA Contact Sites. Journal of Cell Biology, 136(3):501–513.

Slupphaug, G., OIsen, L. C., Helland, D., Aasland, R., and Krokan, H. E. (1991). Cell cycle regulation and in vitro hybrid arrest analysis of the major human uracil-DNA glycosylase. Nucleic Acids Research, 19(19):5131–5137.

Spirtes, P., Glymour, C., and Scheines, R. (2001). Causation, Prediction, and Search. The MIT Press.

Steuerman, Y., Cohen, M., Peshes-Yaloz, N., Valadarsky, L., Cohn, O., David, E., Frishberg, A., Mayo, L., Bacharach, E., Amit, I., et al. (2018). Dissection of influenza infection in vivo by single-cell rna sequencing. Cell systems, 6(6):679–691.

Stuart, T., Butler, A., Hoffman, P., Hafemeister, C., Papalexi, E., III, W. M. M., Hao, Y., Stoeckius, M., Smibert, P., and Satija, R. (2019). Comprehensive integration of single-cell data. Cell, 177:1888–1902.

Su-Feher, L., Rubin, A. N., Silberberg, S. N., Catta-Preta, R., Lim, K. J., Ypsilanti, A. R., Zdilar, I., McGinnis, C. S., McKinsey, G. L., Rubino Jr, T. E., et al. (2022). Single cell enhancer activity distinguishes gabaergic and cholinergic lineages in embryonic mouse basal ganglia. Proceedings of the National Academy of Sciences, 119(15):e2108760119.

Takeuchi, A., Takahashi, Y., Iida, K., Hosokawa, M., Irie, K., Ito, M., Brown, J. B., Ohno, K., Nakashima, K., and Hagiwara, M. (2020). Identification of qk as a glial precursor cell marker that governs the fate specification of neural stem cells to a glial cell lineage. Stem Cell Reports, 15(4):883–897.

Tirosh, I., Izar, B., Prakadan, S. M., Wadsworth, M. H., Treacy, D., Trombetta, J. J., Rotem, A., Rodman, C., Lian, C., Murphy, G., et al. (2016). Dissecting the multicellular ecosystem of metastatic melanoma by single-cell rna-seq. Science, 352(6282):189–196.

Tremblay, R., Lee, S., and Rudy, B. (2016). Gabaergic interneurons in the neocortex: From cellular properties to circuits. Neuron, 91(2):260–292.

Trimm, E. and Red-Horse, K. (2023). Vascular endothelial cell development and diversity. Nature Reviews Cardiology, 20(3):197–210.

Tsunekawa, Y., Kikkawa, T., and Osumi, N. (2014). Asymmetric inheritance of cyclin d2 maintains proliferative neural stem/progenitor cells: A critical event in brain development and evolution. Development, Growth & Differentiation, 56(5):349–357.

Uhlén, M., Fagerberg, L., Hallström, B. M., Lindskog, C., Oksvold, P., Mardinoglu, A., Sivertsson, Å., Kampf, C., Sjöstedt, E., Asplund, A., et al. (2015). Tissue-based map of the human proteome. Science, 347(6220):1260419.

Uhlén, M., Zhang, C., Lee, S., Sjöstedt, E., Fagerberg, L., Bidkhori, G., Benfeitas, R., Arif, M., Liu, Z., Edfors, F., et al. (2017). A pathology atlas of the human cancer transcriptome. Science, 357(6352):eaan2507.

VanderWeele, T. J. and Knol, M. J. (2014). A tutorial on interaction. Epidemiologic Methods, 3(1):33–72.

Watkinson, J., Liang, K.-c., Wang, X., Zheng, T., and Anastassiou, D. (2009). Inference of regulatory gene interactions from expression data using three-way mutual information. Annals of the New York Academy of Sciences, 1158(1):302–313.

Winkler, C. and Yao, S. (2014). The midkine family of growth factors: diverse roles in nervous system formation and maintenance. British Journal of Pharmacology, 171(4):905–912.

Wolf, F. A., Angerer, P., and Theis, F. J. (2018). Scanpy: large-scale single-cell gene expression data analysis. Genome Biology, 19(1):15.

Wolock, S. L., Lopez, R., and Klein, A. M. (2019). Scrublet: Computational identification of cell doublets in single-cell transcriptomic data. Cell Systems, 8(4):281–291.e9.

Wu, T., Hu, E., Xu, S., Chen, M., Guo, P., Dai, Z., Feng, T., Zhou, L., Tang, W., Zhan, L., Fu, X., Liu, S., Bo, X., and Yu, G. (2021). clusterprofiler 4.0: A universal enrichment tool for interpreting omics data. The Innovation, 2(3).

Xia, B. and Yanai, I. (2019). A periodic table of cell types. Development, 146(12):dev169854.

Yang, L., Li, Z., Liu, G., Li, X., and Yang, Z. (2022). Developmental origins of human cortical oligodendrocytes and astrocytes. Neuroscience Bulletin, 38(1):47–68.

Yu, G., Wang, L.-G., Han, Y., and He, Q.-Y. (2012). clusterprofiler: an r package for comparing biological themes among gene clusters. OMICS: A Journal of Integrative Biology, 16(5):284–287. PMID: 22455463.

Yuzwa, S. A., Borrett, M. J., Innes, B. T., Voronova, A., Ketela, T., Kaplan, D. R., Bader, G. D., and Miller, F. D. (2017). Developmental emergence of adult neural stem cells as revealed by single-cell transcriptional profiling. Cell Reports, 21(13):3970–3986.

Zeisel, A., Hochgerner, H., Lönnerberg, P., Johnsson, A., Memic, F., Van Der Zwan, J., Häring, M., Braun, E., Borm, L. E., La Manno, G., et al. (2018). Molecular architecture of the mouse nervous system. Cell, 174(4):999–1014.e22.

Zhang, Z., Wei, S., Du, H., Su, Z., Wen, Y., Shang, Z., Song, X., Xu, Z., You, Y., and Yang, Z. (2019). Zfhx3 is required for the differentiation of late born d1-type medium spiny neurons. Experimental Neurology, 322:113055.

Zheng, K., Huang, H., Yang, J., and Qiu, M. (2022). Origin, molecular specification, and stemness of astrocytes. Developmental Neurobiology, 82(2):149–159.

